# Dose-dependent NFI regulation of progenitor lifespan and output underlying human neocortical malformation

**DOI:** 10.64898/2025.12.27.696675

**Authors:** Qiangqiang Zhang, Guohua Yuan, Elena Albizzati, Jiajun Yang, Zhe Zhao, Xiangyu Yu, Xuyao Chang, Choong Heon Lee, Heng Du, Zhimin Lao, Anjana Krishnamurthy, Xiuli Zhang, Xiaohui Lv, Xing Tang, Shuhan Hu, Yudan Chi, Jian Ma, Richard M. Gronostajski, Linda J. Richards, Jiangyang Zhang, Alexandra L. Joyner, Jason Tchieu, Yinqing Li, Song-Hai Shi

## Abstract

Nuclear Factor I (NFI) misexpressions in humans are associated with severe brain malformations, yet the underlying mechanisms remain poorly understood. Here we show that NFIs regulate the broad lineage progression and lifespan of radial glial progenitors (RGPs), and consequently neocortical development in a dose-dependent manner. Human cerebral organoids carrying patient-mimicking *NFI* mutations exhibit expression level-dependent bidirectional impairments in RGP temporal development, coinciding with patient phenotypes. In mouse models, selective removal of NFIs leads to a dramatic protraction of RGP lineage progression and lifespan, excessive progeny output, and neocortical overgrowth and abnormal folding, whereas overexpression of NFIs accelerates RGP lineage progression, resulting in developmental stage-dependent precocious productions of diverse neural progenies. Moreover, NFIs exhibit a positive auto-regulation and progressive increase in expression, and regulate distinct temporal-specific targets underlying RGP lineage progression. These results suggest that NFIs act as evolutionarily conserved key global temporal regulators of RGP lineage progression and neocortical development.

**HIGHLIGHTS:**

- NFIs affect human RGP temporal development coinciding with patient phenotypes.
- NFI removal protracts RGP lineage progression and lifespan with excessive progeny output.
- NFI overexpression accelerates RGP lineage progression with precocious progeny output.
- NFIs regulate distinct temporal-specific targets underlying RGP lineage progression.

## INTRODUCTION

The neocortex is one of the most complex parts of the brain, consisting of an extraordinarily large number of diverse neurons and glial cells. The immense number and diversity of neural cells is generated by neural stem or progenitor cells, which transit through a developmental program of proliferation, neurogenesis, and gliogenesis, and finally exit the cell cycle and become depleted upon committing to terminal differentiation ^1,2^. The proper lineage progression and lifespan of neural progenitors essentially determine the number and diversity of neural cells in the nervous system. Disruptions in the intricate process of lineage progression and lifespan of neural progenitors can lead to brain malformations, which include various structural abnormalities such as microcephaly and macrocephaly, contributing to developmental delay and intellectual disability. Notably, accumulating evidence has implicated the Nuclear Factor I (NFI) family of transcription factors (TFs) in human brain malformations ^3–12^. Three NFI family members (NFIA, NFIB, NFIX) are abundantly expressed in the developing central nervous system ^13–17^. *NFI* deficiency is linked to head circumference enlargement and polymicrogyria ^3,4,6,18^, whereas *NFI* duplication is associated with small head circumference ^11^. In addition, various *NFI* nonsense or frameshift mutations have been found to be associated with brain abnormalities ^7,8,10,19,20^. While the genetic evidence is compelling, the mechanisms linking NFI misexpressions to brain malformations remain poorly understood.

Radial glial progenitors (RGPs) in the ventricular zone (VZ) of the developing neocortex represent the major neural progenitors responsible for producing diverse neurons and glial cells to constitute the future neocortex ^21–29^. They undergo an organized temporal program of proliferation and differentiation ^23,30–33^. Initially (prior to embryonic day 12, E12, in mice), RGPs largely undergo symmetric proliferative division to amplify the progenitor pool. They then transition to a neurogenic phase during which they predominantly divide asymmetrically to self-renew and, simultaneously, produce deep and superficial layer neurons either directly or indirectly via intermediate progenitors (IPs) in a sequential manner. At the completion of neurogenesis (around E16-E17 in mice), a fraction of RGPs proceeds to gliogenesis to generate astrocytes and oligodendrocytes, while the majority exit the cell cycle upon terminal neurogenic division. Gliogenic RGPs continue to be depleted during the process of gliogenesis, with a small fraction transitioning to adult neural stem cells in the subventricular zone ^34,35^. Notably, this highly organized temporal program of RGP lineage progression occurs at the single cell level ^23,30^, indicating that the temporal state and lifespan of RGPs is well-defined and tightly regulated.

In principle, the lineage progression and lifespan of RGPs could be regulated by both cell intrinsic and extrinsic mechanisms ^36^. It has previously been shown that RGPs display a similar pattern of lineage progression when cultured *in vitro* in isolation and *in vivo* ^37–39^, indicating that the orderly lineage progression and lifespan is a fundamental feature of RGPs and orchestrated by cell intrinsic mechanisms. Nonetheless, the environment also influences RGP lineage progression and progeny output ^40^. Previous studies in the mouse neocortex and retina suggest that some intrinsic factors like orthologs of the Drosophila temporal identity TFs have a conserved function in mammalian neural progenitors ^41–43^. Moreover, progressive transcriptomic changes have also been found to coincide with neocortical RGP temporal development ^44–48^.

The role of NFIs as glial-determining factors in the developing spinal cord has been well documented, with NFIA and NFIB being both necessary and sufficient to promote glial-fate specification in embryonic spinal cord progenitors ^49–52^. Recently, NFIA and NFIB have also been shown to be required for the efficient generation of late-born spinal cord neurons ^17^. Similarly, in the retina, NFIs are required for the specification of Müller glia, the main glial cell of the retina, and bipolar cells, a late-born neuronal subtype ^15^. NFIs are also crucial for midline glia generation and corpus collosum development in the developing forebrain ^53^. In addition, NFIs have been found to regulate neocortical development in seemingly opposite manners; *Nfia* or *Nfib* knockout mice displayed a decrease in differentiated progenies and neocortical volume ^54–56^, whereas *Nfix* knockout mice exhibited an increase in neocortical volume ^57^. These findings in conjunction with human genetic data emphasize the importance of NFIs in regulating neocortical development; yet, the precise roles of NFIs in the developing neocortex require further investigation, especially in the context of brain malformations.

To address this, we integrated human cerebral organoid models and mouse models, and performed systematic analyses of NFI functions in neocortical development. We found that in human cerebral organoids carrying patient-mimicking *NFI* mutations, RGPs exhibit bidirectional impairments in temporal development, coinciding with patient phenotypes. Furthermore, in spatiotemporally specific loss- and gain-of-function studies in mouse models, we showed that NFIs act as global temporal regulators to tune the broad RGP lineage progression rate from proliferation to neurogenesis to gliogenesis to depletion, and consequently RGP lifespan and progeny output, distinct from the previous understanding of NFI primarily as a gliogenic or late-born progeny determinant. Furthermore, by developing and applying integrated single cell multi-omics analyses, we revealed that progressive expression of NFIs in RGPs regulate distinct temporal-specific target TFs coupled with RGP lineage progression and temporal development. These results point to an evolutionarily conserved mechanism of NFIs as global temporal regulators in controlling RGP lineage progression, lifespan, progeny output in a dose-dependent manner under both normal and disease conditions.

## RESULTS

### NFIA/X bidirectionally tune human RGP lineage progression coinciding with patient phenotypes

To investigate the mechanisms underlying human patient brain malformations associated with *NFI* misexpressions, we generated human cerebral organoids and examined their development upon manipulating NFI expression levels to mimic patient conditions **(Figure 1A)**. We focused on NFIA and NFIX, as previous studies indicate that NFIA and NFIB share similar biological functions ^54^. Human embryonic stem cell (hESC) lines carrying loss of function alleles for both *NFIA* and *NFIX* were induced to differentiate into cerebral organoids ^58,59^ **(Figure S1A)**. Interestingly, compared with wild-type (WT) cerebral organoids, *NFIA/X* double mutant (*NFIA;NFIX* DKO) cerebral organoids were substantially larger in size **(Figures 1B and 1C)**, coinciding with the macrocephaly phenotype observed in human patients with *NFI* deficiency ^3,4,7,8,60,61^. We then stained the sections of WT and *NFIA;NFIX* DKO organoids with an antibody against PAX6, a well-established RGP marker or TBR2, a well-established IP marker. We found that the total area and number of PAX6^+^ RGPs were greatly increased **(Figures 1D-1G)**, whereas the relative fraction of TBR2^+^ IPs was significantly reduced **(Figures 1F, 1H, and 1I)**. Consistent with an increase in RGPs, the number of cells expressing brain lipid-binding protein (BLBP), another well-characterized RGP marker, was significantly increased **(Figure S1B)**. These results suggest that loss of NFIA and NFIX leads to an increase in RGP proliferation and a concurrent decrease in IP generation towards differentiation in human cerebral organoids.

**Figure 1.**
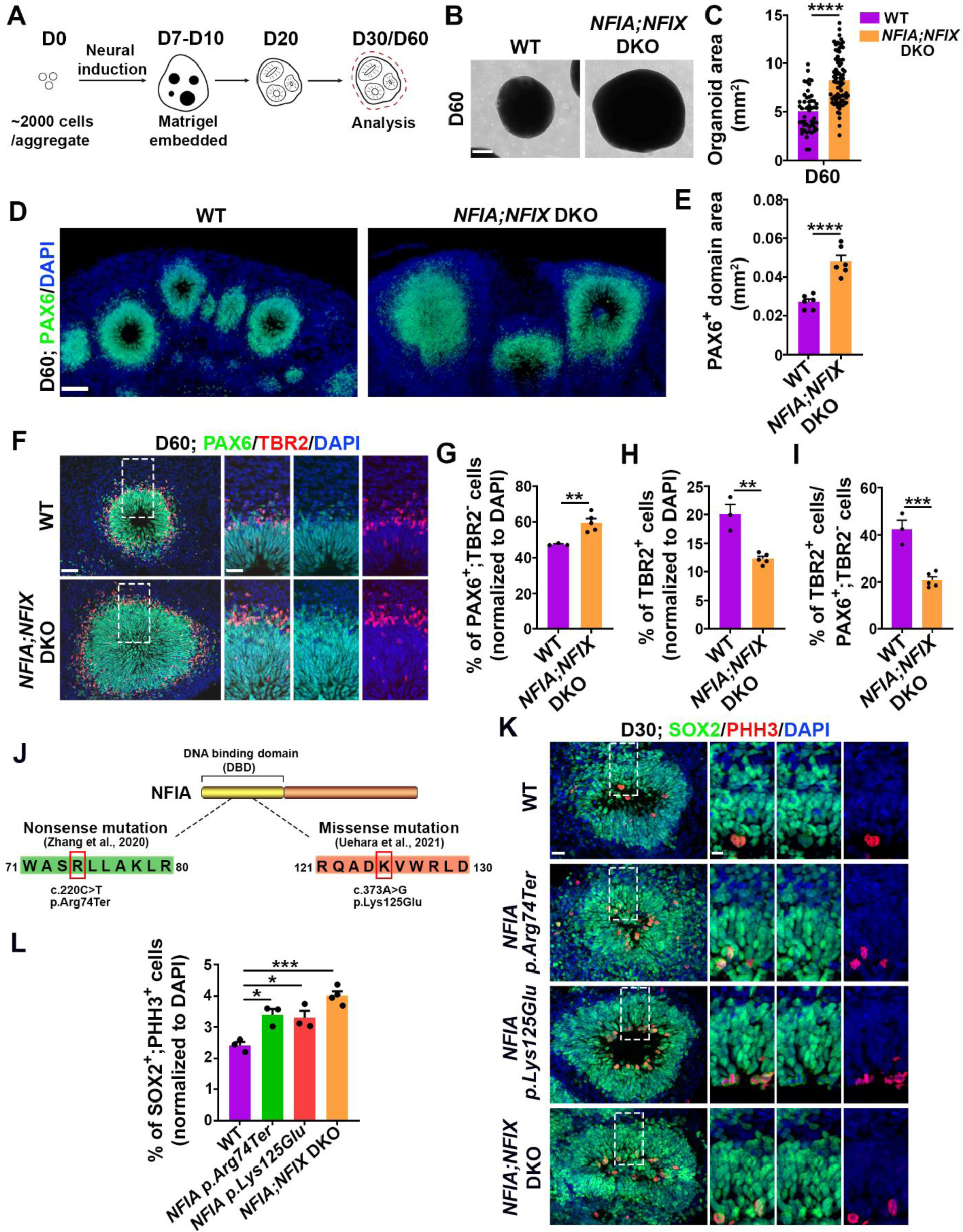
The removal of NFIA and NFIX expression leads to an increase in RGP proliferation and a concurrent decrease in IP generation towards differentiation in human cerebral organoids. **(A)** Schematic of human cerebral organoid generation. **(B)** Representative images of day (D) 60 WT and *NFIA;NFIX* DKO human cerebral organoids. Scale bar, 250 µm. **(C)** Quantification of the 2D surface area of D60 WT and *NFIA;NFIX* DKO human cerebral organoids. **(D)** Representative images of D60 WT and *NFIA;NFIX* DKO human cerebral organoid sections stained for PAX6 (green) and with DAPI (blue). Scale bar, 100 µm. **(E)** Quantification of the area of the PAX6^+^ domain (WT, n = 6 organoids; *NFIA;NFIX* DKO, n = 6 organoids). **(F)** Representative images of D60 WT and *NFIA;NFIX* DKO human cerebral organoid sections stained for PAX6 (green) and TBR2 (red), and with DAPI (blue). Scale bars: 50 µm (left) and 25 µm (right). **(G-I)** Quantifications of the percentages of PAX6^+^;TBR2^-^ (RGP) **(G)** and TBR2^+^ (IP) **(H)** cells out of the total cells, and the ratio of TBR2^+^/PAX6^+^;TBR2^-^ cells **(I)** in D60 WT and *NFIA;NFIX* DKO human cerebral organoid sections (WT, n = 3 organoids; *NFIA;NFIX* DKO, n = 5 organoids). **(J)** Schematic of the DNA binding domain (DBD) of the human NFIA protein, indicating the locations of two pathogenic variants: the p.Arg74Ter nonsense mutation and the p.Lys125Glu missense mutation. **(K)** Representative images of D30 WT, *NFIA p.Arg74Ter*, *NFIA p.Lys125Glu*, and *NFIA;NFIX* DKO human cerebral organoid sections stained for SOX2 (green) and PHH3 (red), and with DAPI (blue). Scale bars: 25 µm (left) and 10 µm (right). **(L)** Quantifications of the percentages of SOX2^+^;PHH3^+^ cells out of the total cells in D30 WT, *NFIA p.Arg74Ter*, *NFIA p.Lys125Glu*, and *NFIA;NFIX* DKO human cerebral organoid sections (bottom, WT, n = 3 organoids; *NFIA p.Arg74Ter*, n = 3 organoids; *NFIA p.Lys125Glu*, n = 3 organoids *NFIA;NFIX* DKO, n = 4 organoids). Data are presented as mean ± SEM. *, P < 0.05; **, P < 0.01; ***, P < 0.001; ****, P < 0.0001. Statistical analysis was performed using unpaired Student’s t-test **(C, E, G-I)** and One-way ANOVA **(L)**.

To further explore the link between *NFI* mutations and human brain malformation, we generated two additional types of human cerebral organoids carrying two pathogenic variants of the *NFI* gene – the *NFIA p.Arg74Ter* variant, which is a *de novo* heterozygous nonsense mutation and causes macrocephaly in human patients ^19^, and the *NFIA p.Lys125Glu* variant, which is a recurrent heterozygous missense mutation in the DNA binding domain of NFIA, representing a loss-of-function pathogenic allele and resulting in macrocephaly in human patients ^20^ **(Figure 1J)**. Notably, both types of human cerebral organoids carrying the pathogenic *NFI* gene variants exhibited a significant increase in RGP proliferation compared with WT control organoids **(Figures 1K and 1L)**, in line with the macrocephaly phenotype observed in human patients carrying the corresponding pathogenic variants ^19,20^.

We next investigated the effects of NFIA and NFIX overexpression on human RGP lineage progression. We engineered hESC lines harboring doxycycline-inducible expression of NFIA and NFIX. Strikingly, NFIA and NFIX overexpressing (*NFIA/X* OE) cerebral organoids were substantially smaller in size than WT control organoids **(Figures 2A and 2B)**, consistent with the microcephaly phenotype observed in human patients with NFI microduplication ^11,62^. Moreover, the total area of the PAX6^+^ domain was greatly reduced **(Figures 2C and 2D)**, indicating a premature loss of RGPs. In addition, we observed a substantial increase in the relative fraction of TBR2^+^ IPs **(Figures 2E-2H)**, suggesting a precocious increase in differentiation. These results indicate that NFIA and NFIX overexpression results in a premature loss of RGPs and a concomitant precocious differentiation in human cortical organoids.

**Figure 2.**
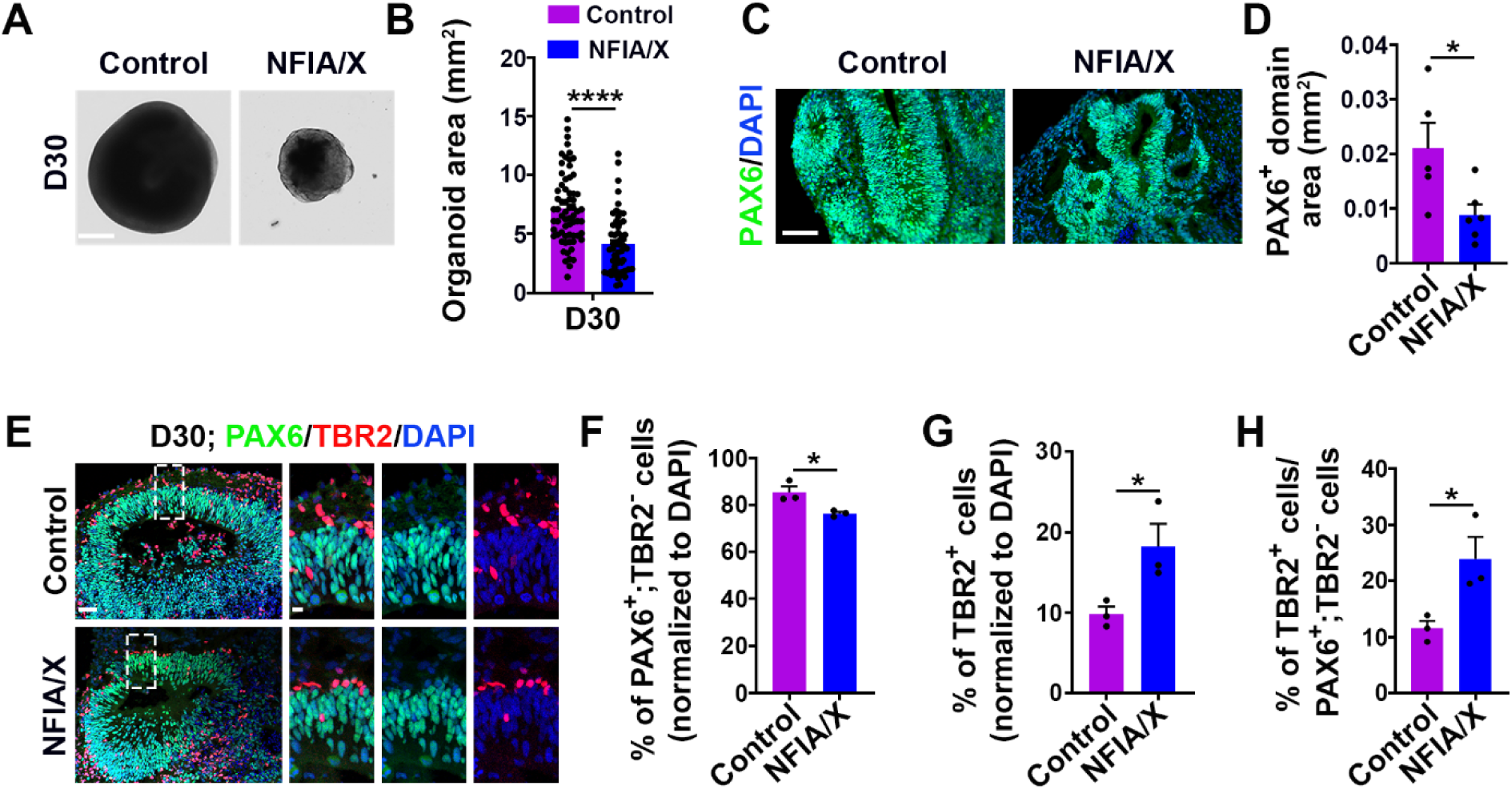
NFIA and NFIX overexpression results in a premature loss of RGPs and a concomitant precocious differentiation in human cerebral organoids. **(A)** Representative images of D30 WT and *NFIA;NFIX* DOE (NFIA/X) human cerebral organoids. Scale bar, 250 µm. **(B)** Quantification of the 2D surface area of D30 WT and NFIA/X human cerebral organoids. **(C)** Representative images of D30 WT and NFIA/X human organoid sections stained for PAX6 (green) and with DAPI (blue). Scale bar, 100 µm. **(D)** Quantification of the area of the PAX6^+^ domain in D30 WT and NFIA/X human cerebral organoids. (WT, n = 5 organoids; NFIA/X, n = 5 organoids). **(E)** Representative images of D30 WT and NFIA/X human cerebral organoid sections stained for PAX6 (green) and TBR2 (red), and with DAPI (blue). Scale bars, 100 µm (left) and 25 µm (right). **(F-H)** Quantifications of the percentages of PAX6^+^;TBR2^-^ (RGP) **(F)** or TBR2^+^ (IP) **(G)** cells out of the total cells, and the ratio of TBR2^+^/PAX6^+^;TBR2^-^ cells **(H)** in D30 WT and NFIA/X human organoid sections (WT, n = 3 organoids; NFIA/X, n = 3 organoids). Data are presented as mean ± SEM. *, P < 0.05; ****, P < 0.0001. Statistical analysis was performed using unpaired Student’s t-test.

Taken together, these findings demonstrate that in parallel with patient phenotypes, abnormal NFI expression or dosage bidirectionally affects human RGP lineage progression and temporal development in proliferation and differentiation.

### NFIA/X removal in RGPs leads to an excessively enlarged cortex with extensive folding

To further examine the bidirectional role of NFIs in regulating RGP lineage progression and temporal development in the developing neocortex, we next performed in-depth analyses of NFI function in RGPs in mice, which allows for full lineage progression and cortical development assessment. We took advantage of the conditional mutant mice of *Nfia*, *Nfib*, and *Nfix* (i.e., *Nfia^fl/fl^*, *Nfib^fl/fl^*, and *Nfix^fl/fl^*) ^57,63,64^, and crossed them with *Emx1-Cre* mice ^65^, in which Cre recombinase is mostly expressed in cortical RGPs starting at ∼E9.5. *Emx1-Cre;Nfia^fl/fl^;Nfib^fl/fl^;Nfix^fl/fl^* conditional triple knockout mice died around the neonatal stage, precluding a full assessment of RGP lineage progression and cortical development. To circumvent this, we focused our analysis on *Emx1-Cre;Nfia^fl/fl^;Nfix^fl/fl^* conditional double knockout mice (hereinafter referred to as *Nfia;Nfix* cDKO). *Nfia;Nfix* cDKO mice were born at the expected frequency and survived to adulthood with a clear loss of NFIA and NFIX expression in the cortex starting from the embryonic stage **(Figure S2A)**. The brain of *Nfia;Nfix* cDKO mice was significantly larger than that of WT littermate control mice at postnatal day (P) 21 **(Figures 3A and 3B)**. Magnetic resonance imaging (MRI) analysis confirmed that the cortex was substantially enlarged across the entire rostrocaudal axis, especially in the medial region **(Figures 3C and 3D)**.

**Figure 3.**
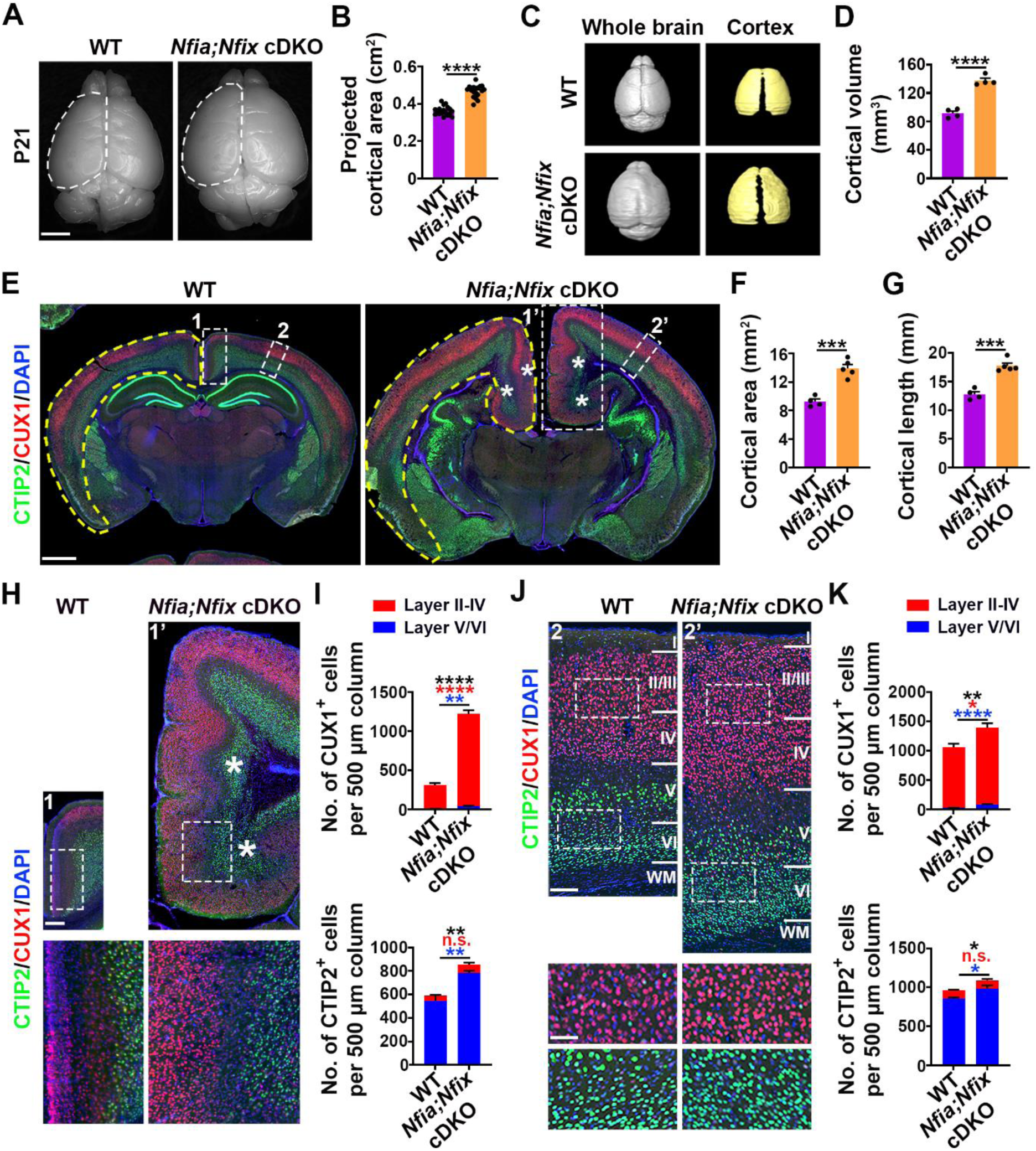
The removal of NFIA and NFIX in RGPs leads to an excessively enlarged cortex with extensive folding. **(A)** Representative whole-mount images of P21 wild-type (WT) and *Nfia;Nfix* cDKO brains. Broken lines indicate the cerebral hemisphere. Scale bar, 0.25 cm. **(B)** Quantification of the projected cortical area (WT, n = 18 brains; *Nfia;Nfix* cDKO, n = 18 brains). **(C)** MRI images of P21 WT and *Nfia;Nfix* cDKO brains. **(D)** Quantification of P21 WT and *Nfia;Nfix* cDKO cortical volume (WT, n = 4 brains; *Nfia;Nfix* cDKO, n = 4 brains). **(E)** Representative images of P21 WT and *Nfia;Nfix* cDKO brain sections stained for CTIP2 (green) and CUX1 (red), and with DAPI (blue). Yellow dashed outlines delineate the total cortical area. White asterisks indicate the dramatic cortical folding in the medial region. Scale bar, 1 mm. (**F and G)** Quantifications of the cortical area **(F)** and length **(G)** (WT, n = 4 brains; *Nfia;Nfix* cDKO, n = 5 brains). **(H)** Representative images of the medial region of P21 WT and *Nfia;Nfix* cDKO cortices stained for CTIP2 (green) and CUX1 (red), and with DAPI (blue). White asterisks indicate the extensive cortical folding. Scale bars, 250 μm (top) and 100 μm (bottom). **(I)** Quantifications of the number of CUX1^+^ (top) and CTIP2^+^ (bottom) neurons per 500 μm width area in the medial region (n = 4 brains for each genotype). **(J)** Representative images of the dorsal region of P21 WT and *Nfia;Nfix* cDKO cortices stained for CTIP2 (green) and CUX1 (red), and with DAPI (blue). Scale bars, 100 μm (top) and 50 μm (bottom). **(K)** Quantifications of the number of CUX1^+^ (top) and CTIP2^+^ (bottom) neurons per 500 μm width area in the dorsal region (n = 3 brains for each genotype). Data are presented as mean ± SEM. *, P < 0.05; **, P < 0.01; ***, P < 0.001; ****, P < 0.0001; n.s., not significant. Statistical analysis was performed using unpaired Student’s t-test.

The enlarged cortex in the *Nfia;Nfix* cDKO brain indicates abnormalities in neuronal production. To examine this, we stained P21 brain sections with antibodies against CTIP2 (green), a layer V/VI neuronal marker, and CUX1 (red), a layer II-IV neuronal marker ^66^. Consistent with the cortical enlargement, we observed a marked increase in the overall length and area of the whole cortex with normal lamination in the *Nfia;Nfix* cDKO brain compared with the WT brain **(Figures 3E-3G and S2B)**. Furthermore, in the medial region of the *Nfia;Nfix* cDKO cortex exhibiting the most substantial volume increase, we observed consistent and extensive enlargement and folding **(Figures 3E and 3H right, asterisks)**, which never occurred in the WT cortex **(Figures 3E and 3H left)**. Both the densities and total numbers of CTIP2^+^ deep layer and CUX1^+^ superficial layer neurons were dramatically increased in the medial and dorsal regions of the *Nfia;Nfix* cDKO cortex compared with the WT cortex **(Figures 3H-3K)**. In alignment with the folding, the extent of neuronal production increase in the medial region was more substantial than that in the dorsolateral region. We also observed extensive enlargement and folding of the hippocampus in the *Nfia;Nfix* cDKO brain **(Figure S2B)**. The densities of glial cells did not show any obvious change **(Figures S2C, S2D, S2F and S2G)**; however, because of the large increase in the total area and volume of the *Nfia;Nfix* cDKO cortex, the net production of glial cells was also substantially enhanced **(Figures S2E and S2H)**.

Given that NFIA and NFIX are expressed in some postmitotic cortical neurons ^13^, we tested the possibility that NFIA and NFIX may function in postmitotic neurons to contribute to the cortical expansion and folding phenotype observed in *Nfia;Nfix* cDKO mice. We crossed *Nfia^fl/fl^;Nfix^fl/fl^* mice with the *Nex-Cre* mouse line ^67^, in which Cre recombinase is selectively expressed in postmitotic cortical neurons. Compared with the WT littermate control, we observed no obvious change in the overall area or length of the mutant cortex, or in the total numbers of CUX1^+^ superficial or CTIP2^+^ deep layer neurons in the *Nex-Cre;Nfia^fl/fl^;Nfix^fl/fl^*cDKO cortex at P21 **(Figures S2I-S2M)**, indicating that the extensive enlargement and folding observed in the *Nfia;Nfix* cDKO cortex is unlikely due to the loss of NFIA and NFIX in postmitotic neurons. Taken together, these results suggest the selective removal of NFIA and NFIX in RGPs leads to a dramatic increase in the production of both deep and superficial layer neurons as well as glial cells, and consequently the extensive enlargement and folding of the cortex, especially in the medial region.

### NFIA/X removal prolongs RGP lineage progression and lifespan

The excessive neural progeny production and enlargement of the *Nfia;Nfix* cDKO cortex indicate that the behavior and lineage progression of RGPs are altered due to the loss of NFIA and NFIX. To test this, we systematically examined the temporal development and lifespan of cortical RGPs. We stained brain sections of WT and *Nfia;Nfix* cDKO cortices at different developmental stages with an antibody against RGP marker PAX6 **(Figure 4A)**. Notably, the PAX6^+^ domain in the *Nfia;Nfix* cDKO cortex was significantly increased at E13.5 compared with the WT cortex **(Figures 4A and 4B)**. This increase became more pronounced at E15.5, E17.5, and P1 **(Figures 4A, 4B, S3A and S3B)**. As development proceeded, the PAX6^+^ RGPs in the dorsal region of the WT cortex progressively diminished and became largely undetectable by P14, whereas the PAX6^+^ RGPs in the dorsal region of the *Nfia;Nfix* cDKO cortex remained abundant at P60 **(Figures 4A and 4B)** or even later at 6 months (**Figure S3C**). These long-lived PAX6^+^ RGPs possessed radial glial fibers, as revealed by RC2 staining, and retained the ability to divide, as reflected by 5-Bromo-2’-deoxyuridine (BrdU) incorporation **(Figures 4C and 4D)**. These results suggest that the removal of NFIA and NFIX leads to a drastic change in lineage progression and lifespan of RGPs in the developing cortex.

**Figure 4.**
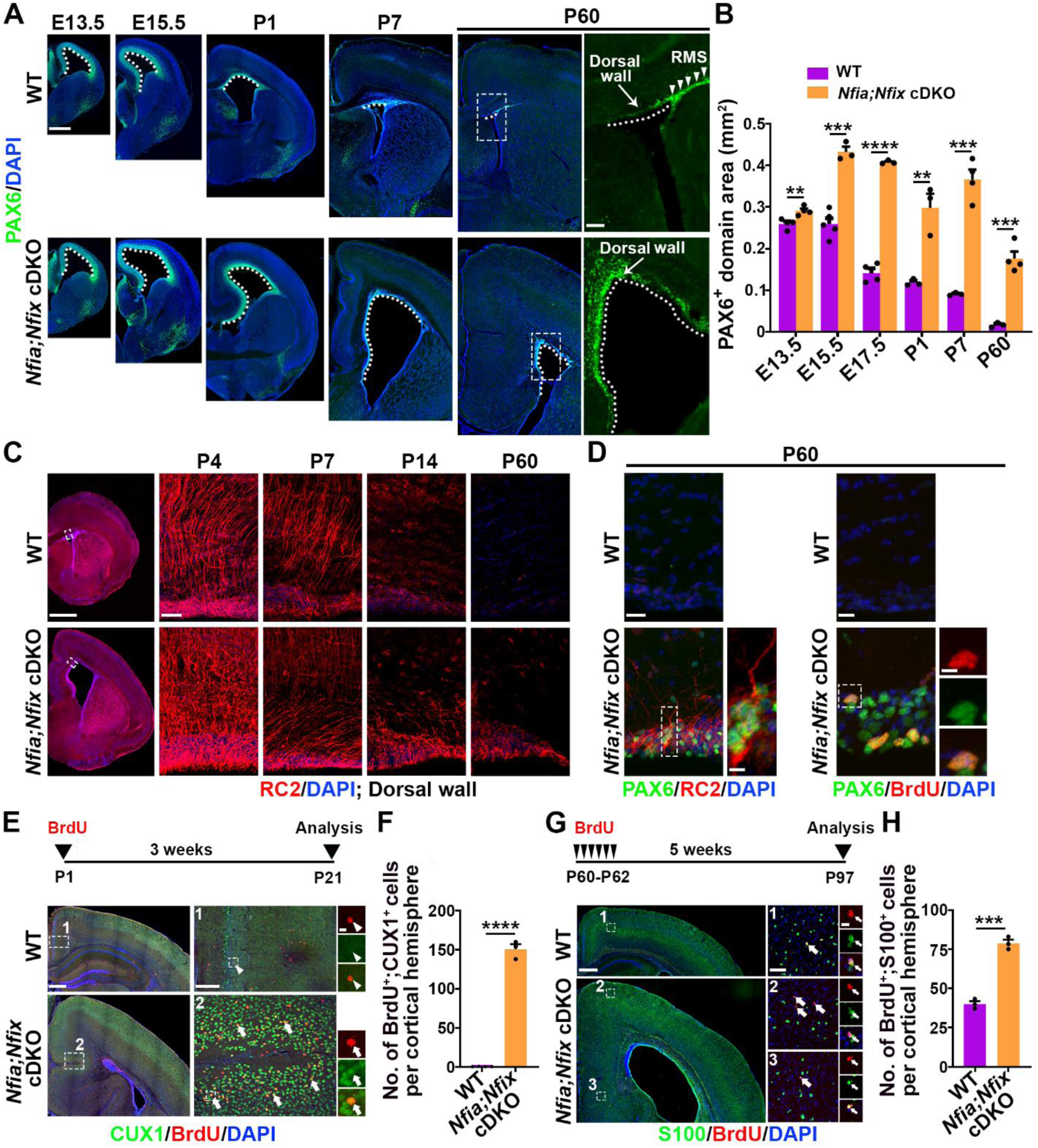
The removal of NFIA and NFIX dramatically prolongs RGP lifespan. **(A)** Representative images of E13.5, E15.5, P1, P7, and P60 WT and *Nfia;Nfix* cDKO cortices stained for PAX6 (green), an RGP marker, and with DAPI (blue). The dashed lines indicate the boundaries of the dorsal wall of the lateral ventricle. Arrows indicate the extensive presence of PAX6^+^ RGPs in the dorsal wall of the lateral ventricle in the *Nfia;Nfix* cDKO cortex, even at P60, compared with the WT control. RMS, rostral migratory stream. Scale bars, 500 µm (left) and 100 µm (right). **(B)** Quantification of the area of the PAX6^+^ domain in the dorsal wall (E13.5: WT, n = 4 brains; *Nfia;Nfix* cDKO, n = 4 brains; E15.5: WT, n = 5 brains; *Nfia;Nfix* cDKO, n = 3 brains; E17.5: WT, n = 4 brains; *Nfia;Nfix* cDKO, n = 3 brains; P1: WT, n = 3 brains; *Nfia;Nfix* cDKO, n = 3 brains; P7: WT, n = 3 brains; *Nfia;Nfix* cDKO, n = 4 brains; P60: WT, n = 3 brains; *Nfia;Nfix* cDKO, n = 3 brains). **(C)** Representative images of the dorsal wall of P4, P7, P14, and P60 WT and *Nfia;Nfix* cDKO cortices stained for RC2 (red), an RGP marker, and with DAPI (blue). The white dashed boxes indicate dorsal wall areas showed on the right. Scale bars, 1 mm (left) and 50 µm (right). **(D)** Representative images of the dorsal wall of P60 WT and *Nfia;Nfix* cDKO cortices stained for PAX6 (green) and RC2 (red, left) or BrdU (red, right), and with DAPI (blue). High-magnification images of the white dashed boxes were showed on the right. Scale bars, 20 µm (left top), 5 µm (left bottom), 10 µm (right top), and 5 µm (right bottom). **(E)** (top) Schematic of BrdU birthdating assay. (bottom) Representative images of P21 WT and *Nfia;Nfix* cDKO cortices stained for CUX1 (green) and BrdU (red) administered at P1, and with DAPI (blue). High-magnification images of the medial cortex (broken line rectangles 1 and 2) are shown to the right. The arrowhead indicates a BrdU^+^;CUX1^-^ cell. Arrows indicate BrdU^+^;CUX1^+^ cells. Scale bars, 500 μm (left), 100 μm (middle), and 10 μm (right). **(F)** Quantification of the number of BrdU^+^;CUX1^+^ cells per cortical hemisphere in WT and *Nfia;Nfix* cDKO cortices (WT, n = 4 brains; *Nfia;Nfix* cDKO, n = 3 brains). **(G)** (top) Schematic of BrdU birthdating assay. (bottom) Representative images of P97 WT and *Nfia;Nfix* cDKO cortices stained for S100 (green) and BrdU (red) administered at P60-P62, and with DAPI (blue). High-magnification images of the medial cortex (broken line rectangles 1, 2 and 3) are shown to the right. The arrow indicates a BrdU^+^;S100^+^ cell. Scale bars, 500 μm (left), 50 μm (middle), and 10 μm (right). **(H)** Quantification of the number of BrdU^+^;S100^+^ cells per cortical hemisphere in WT and *Nfia;Nfix* cDKO cortices (WT, n = 3 brains; *Nfia;Nfix* cDKO, n = 3 brains). Data are presented as mean ± SEM. **, P < 0.01; ***, P < 0.001; ****, P < 0.0001. Statistical analysis was performed using unpaired Student’s t-test.

To further assess RGP lineage progression, we examined the temporal developmental program of proliferation, neurogenesis, and gliogenesis. During normal development, RGPs in the developing cortex largely undergo symmetric proliferative division prior to E12.5. At E13.5, RGPs predominantly undergo asymmetric neurogenic division to produce neurons either directly or indirectly via IPs ^23,30,31^. In the *Nfia;Nfix* cDKO cortex, we observed a significant increase in PAX6^+^ RGPs and a concomitant decrease in TBR2^+^ IPs compared with WT at E13.5 **(Figures S3D-S3F)**, suggesting that a significant number of RGPs in the *Nfia;Nfix* cDKO cortex undergo symmetric proliferative division at the expense of asymmetric neurogenic division. Indeed, paired cell analysis showed a significantly higher fraction of RGPs from the E13.5 *Nfia;Nfix* cDKO cortex underwent symmetric proliferative division compared with WT **(Figures S3G and S3H)**. Together, these results indicate that the removal of NFIA and NFIX extends the symmetric proliferation phase of RGPs.

A similar increase in PAX6^+^ RGPs and concurrent decrease in TBR2^+^ IPs in the *Nfia;Nfix* cDKO cortex was also observed at E15.5 and E17.5 **(Figures S3D-S3F)**. Consistent with this, the rate of cell cycle exit in the E14.5 *Nfia;Nfix* cDKO cortex was significantly decreased relative to the WT cortex **(Figures S4A and S4B)**. The number of CTIP2^+^ new-born neurons in the E15.5 *Nfia;Nfix* cDKO cortex was also drastically reduced **(Figures S4C and S4D)**. Given the increased number of deep and superficial layer neurons observed in the *Nfia;Nfix* cDKO cortex at P21, these results suggest that neurogenesis in the *Nfia;Nfix* cDKO cortex was prolonged. Indeed, we found that compared with the WT cortex, neuronal production in the *Nfia;Nfix* cDKO cortex was extended towards the later developmental stages **(Figures 4E and 4F)**. This prolongation and enhancement of neurogenesis was especially prominent in the dorsomedial region of the cortex. We also observed that the densities of glial cells were substantially reduced in the *Nfia;Nfix* cDKO cortex at P1 relative to the WT cortex **(Figures S4E and S4F)**. However, the overall production of glial cells was substantially enhanced in the *Nfia;Nfix* cDKO cortex at later postnatal stages **(Figures 4G, 4H, S4G and S4H)**. Together, these results suggest that the onset of gliogenesis in the *Nfia;Nfix* cDKO cortex is delayed, but once the gliogenesis begins, it also has an extended duration.

### NFIA/X overexpression accelerates RGP lineage progression

To further test the role of NFIA and NFIX as key temporal regulators of RGPs, we examined the effect of NFIA/X overexpression on RGP lineage progression and lifespan. We introduced plasmids expressing EGFP or NFIA and NFIX (NFIA/X) together with EGFP into RGPs in the VZ of the developing neocortex at E12.5 and examined their temporal development at E15.5 **(Figures S4I)**. As expected, control RGPs expressing EGFP underwent lineage development to produce progeny that migrated radially to the subventricular zone, intermediate zone, and cortical plate, while self-renewing to maintain a substantial number of PAX6^+^ RGPs in the VZ. For RGPs overexpressing NFIA/X, the fraction of PAX6^+^ RGPs maintained in the VZ was significantly reduced **(Figures S4I-S4K)**. These results indicate that overexpression of NFIA/X leads to a premature depletion of RGPs, likely due to an accelerated lineage progression and a precocious exit from the cell cycle.

To directly test this, we examined the progeny output of RGPs expressing EGFP or EGFP/NFIA/X at different developmental stages. We introduced plasmids carrying EGFP or EGFP/NFIA/X into RGPs at E12.5, E15.5, or E17.5, and examined their progeny output at P21 **(Figure 5)**. Control RGPs expressing EGFP at E12.5, E15.5, and E17.5 largely generated deep layer neurons, superficial layer neurons, and glial cells with some layer II neurons, respectively **(Figures 5A-5F)**, representing an orderly lineage progression and neural progeny output. In comparison, RGPs expressing EGFP/NFIA/X at E12.5 produced a similar proportion of deep and superficial layer neurons, indicating a premature generation of late-born superficial layer neurons. Consistent with this, we observed a significant increase in the fraction of EGFP/NFIA/X-expressing cells positive for the late-born superficial layer neuron marker CUX1 and a concurrent decrease in the fraction of cells positive for the early-born deep layer neuron marker CTIP2 **(Figures 5A and 5D)**, suggesting accelerated lineage progression **(Figure 5G)**. Similarly, RGPs expressing EGFP/NFIA/X at E15.5 and E17.5 generated upper layer II neurons and glial cells, and glial cells only, respectively **(Figures 5B, 5C, 5E, and 5F)**, corresponding to later-born neural progenies **(Figure 5G)**. Together, these results suggest that overexpression of NFIA/X in RGPs accelerates lineage progression, resulting in precocious productions of neural progeny. Strikingly, the exact neural progeny output depends on the developmental stage. Thus, NFIA and NFIX, as key global temporal regulators, control overall RGP lineage progression and temporal identity, but not the fate specification of any particular cell type(s), in contrast to their previously suggested role in generating glia or late-born neurons ^15,17,49^.

**Figure 5.**
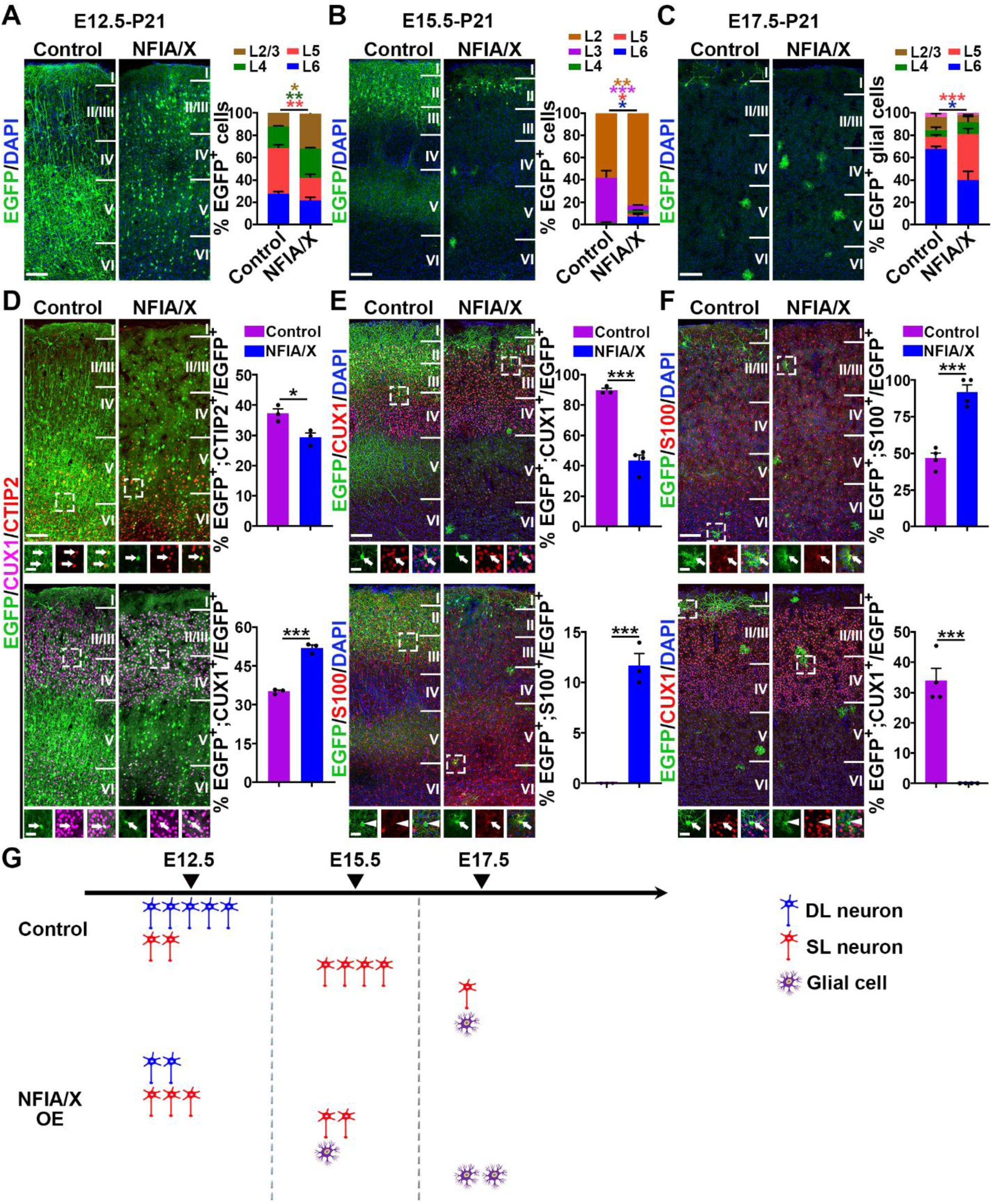
NFIA and NFIX overexpression accelerates RGP lineage progression and neural progeny output. **(A)** (left) Representative images of P21 cortices electroporated with EGFP (green) control or NFIA and NFIX (NFIA/X) overexpression plasmids at E12.5 stained for EGFP (green) and with DAPI (blue). (right) Quantification of the percentage of EGFP^+^ cells in different layers of the cortex (Control, n = 3 brains; NFIA/X, n = 3 brains). Scale bar, 100 µm. **(B)** (left) Representative images of P21 cortices electroporated with EGFP (green) control or NFIA/X overexpression plasmids at E15.5 stained for EGFP (green) and with DAPI (blue). (right) Quantification of the percentage of EGFP^+^ cells in different layers of the cortex (Control, n = 3 brains; NFIA/X, n = 4 brains). Scale bar, 100 µm. **(C)** (left) Representative images of P21 cortices electroporated with EGFP (green) control or NFIA/X overexpression plasmids at E17.5 stained for EGFP (green) and with DAPI (blue). (right) Quantification of the percentage of EGFP^+^ glial cells in different layers of the cortex (Control, n = 4 brains; NFIA/X, n = 4 brains). Scale bar, 100 µm. **(D)** (left) Representative images of P21 cortices electroporated with EGFP (green) control or NFIA/X overexpression plasmids at E12.5 stained for CTIP2 (red) and CUX1 (magenta). High-magnification images (dashed line squares) are shown at the bottom. Arrows indicate EGFP^+^;CTIP2^+^ or EGFP^+^;CUX1^+^ cells. Scale bars, 100 µm (top) and 25 µm (bottom). (right) Quantification of the percentage of EGFP^+^ cells positive for CTIP2 or CUX1 (Control, n = 3 brains; NFIA/X, n = 3 brains). **(E)** (left) Representative images of P21 cortices electroporated with EGFP (green) control or NFIA/X overexpression plasmids at E15.5 stained for CUX1 (red, top) or S100 (red, bottom), and with DAPI (blue). High-magnification images (dashed line squares) are shown at the bottom. Arrows indicate EGFP^+^;CUX1^+^ or EGFP^+^;S100^+^ cells. Arrowheads indicate EGFP^+^;S100^-^ cells. Scale bars, 100 µm (top) and 25 µm (bottom). (right) Quantification of the percentage of EGFP^+^ cells positive for CUX1 or S100 (Control, n = 3 brains; NFIA/X, n = 4 brains). **(F)** (left) Representative images of P21 cortices electroporated with EGFP (green) control or NFIA/X overexpression plasmids at E17.5 stained for S100 (red, top) or CUX1 (red, bottom), and with DAPI (blue). High-magnification images (dashed line squares) are shown at the bottom. Arrows indicate EGFP^+^;S100^+^ or EGFP^+^;CUX1^+^ cells. Arrowheads indicate EGFP^+^;CUX1^-^ cells. Scale bars, 100 µm (top) and 25 µm (bottom). (right) Quantification of the percentage of EGFP^+^ cells positive for S100 or CUX1 (Control, n = 4 brains; NFIA/X, n = 4 brains). **(G)** Schematic summarizing accelerated RGP lineage progression and distinct neural progeny output at different developmental stages when NFIA/X are overexpressed. DL, deep layer; SL, superficial layer. Data are presented as mean ± SEM. *, P < 0.05; **, P < 0.01; ***, P < 0.001. Statistical analysis was performed using unpaired Student’s t-test.

### Progressive expression and activity of NFIs in regulating RGP lineage progression

Having found that NFIA and NFIX regulate RGP lineage progression, lifespan, and progeny output in the developing neocortex, we next examined their expression dynamics and activity in RGPs. The protein expression levels of NFIA and NFIX progressively increase along with RGP temporal development and lineage progression **(Figures 6A and 6B)**. By employing the luciferase reporter assay, a commonly used tool to study gene expression at the transcriptional level, we found that in the presence of NFIA and *Nfia* promoter or cis-regulatory elements (CREs), the luciferase activity was greatly enhanced **(Figure 6C)**, indicating that NFIA positively regulates the *Nfia* transcription mediated by its promoter or CRE. In other words, NFIA is capable of binding to its own promoter and CRE, and exerts a positive auto-regulation of its expression. Similarly, we also found that NFIB and NFIX positively regulated their own expression via their promoters and CREs **(Figures 6D and S5A-S5C)**. This is consistent with the previous observation that NFIB binds to its own enhancer ^68^. These results suggest that NFIs exert a positive auto-regulation in their expression, which supports the progressive increase in their expression in RGPs along with the temporal development.

**Figure 6.**
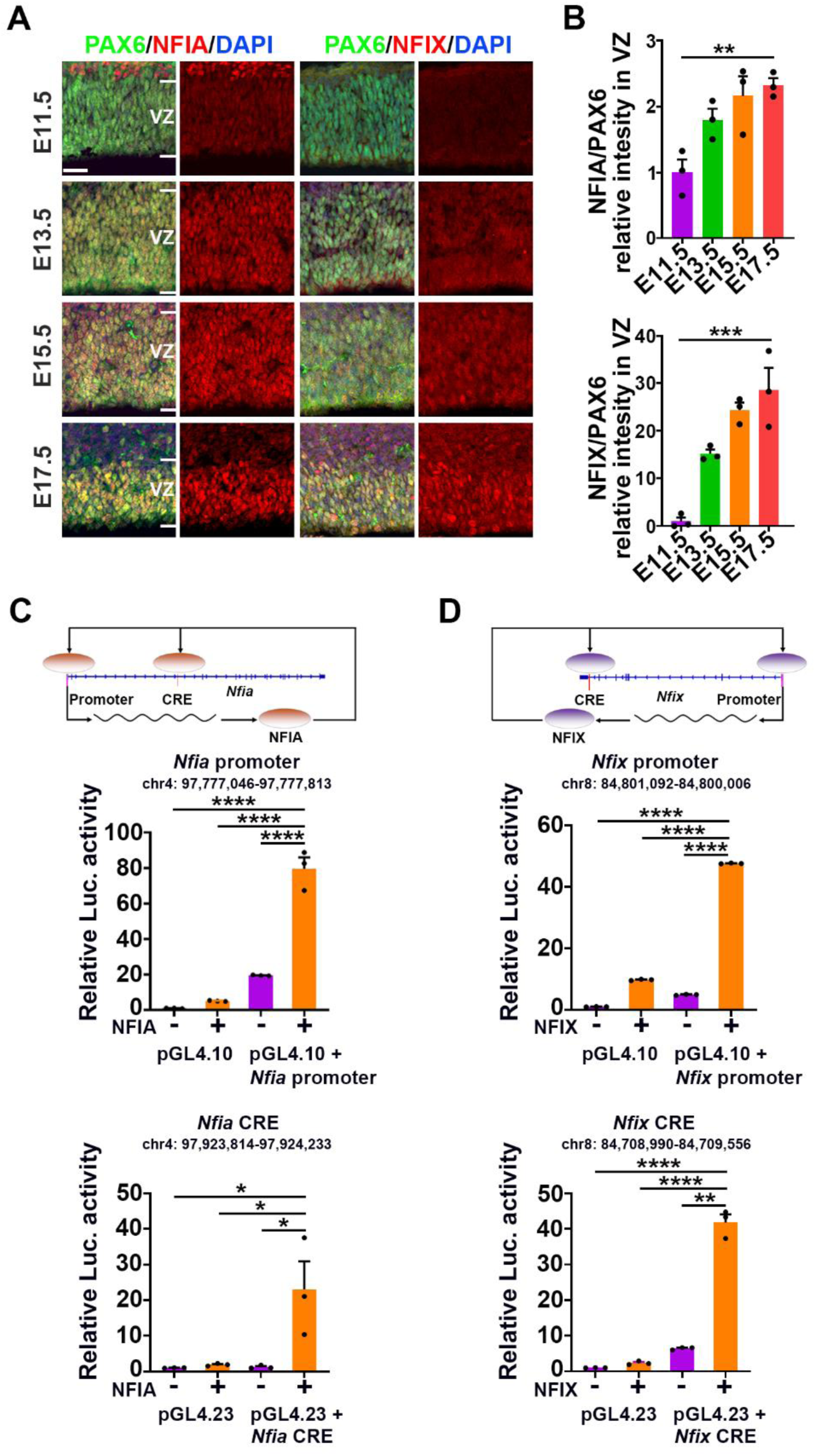
Transcriptional autoregulation controls the progressive increase of NFI expression in RGPs. **(A)** Representative images of the VZ of the developing mouse neocortices at different embryonic stages (E11.5, E13.5, E15.5, and E17.5) stained for PAX6 (green) and either NFIA (red, left), or NFIX (red, right), and with DAPI (blue). Note that the expression of NFIA, or NFIX progressively increases in RGPs as development proceeds. White lines indicate the boundaries of VZ. Scale bar, 25 µm. **(B)** Quantification of the relative intensity of NFIA, or NFIX in RGPs compared to that of PAX6. Fold change of expression was calculated by normalizing to the mean level of expression at E11.5 (n = 3 brains for each time point). **(C)** Luciferase reporter assay was performed to measure activity of NFIA-activated transcription from the *Nfia* promoter and cis-regulatory elements (CREs). The schematic (top) showed that NFIA protein bound the promoter and CRE of *Nfia* gene and amplified its gene transcription by feed-back positive autoregulation. Co-transfection of the NFIA expression construct and the *Nfia* promoter (middle)/*Nfia* CRE (bottom) resulted in a significantly increased level of relative luciferase activity. n=3 biological replicates per group. Luc., luciferase. **(D)** Luciferase reporter assay was performed to measure activity of NFIX-activated transcription from the *Nfix* promoter and CRE. The schematic (top) showed that NFIX protein bound the promoter and CRE of *Nfix* gene and amplified its gene transcription by feed-back positive autoregulation. Co-transfection of the NFIX expression construct and the *Nfix* promoter (middle)/*Nfix* CRE (bottom) resulted in a significantly increased level of relative luciferase activity. n=3 biological replicates per group. Luc., luciferase. Data are presented as mean ± SEM. *, P<0.05; **, P < 0.01; ***, P < 0.001; ****, P<0.0001. Statistical analysis was performed using One-way ANOVA.

Given that NFIs are TFs, the progressive expression of NFIs in RGPs raises the intriguing possibility that NFIs regulate specific targets coupled with RGP lineage progression. To test this, we generated a multi-omics dataset by developing and applying an integrated in-depth single nuclear assay for transposase-accessible chromatin with high throughput (ATAC) sequencing (snATAC-seq) and scRNA-seq (scAnR-seq), as well as comprehensive gene regulation analyses, in the same RGPs across their temporal development (please also see *Yuan et al., bioRxiv 2025*)^69^. This allowed us to systematically examine their regulatory activities by jointly analyzing the expression of *Nfia*, *Nfib*, and *Nfix* as well as the accessibility of DNA regulatory motifs bounded by NFI proteins in the same individual RGPs across time. The mRNA expression level and the DNA binding motif accessibility of *Nfia*/NFIA, *Nfib*/NFIB, and *Nfix*/NFIX exhibited concomitant and progressive increases as RGPs moved through the developmental program **(Figure 7A)**, indicating a persistent and global regulation by NFIs across the entire RGP lineage progression.

**Figure 7.**
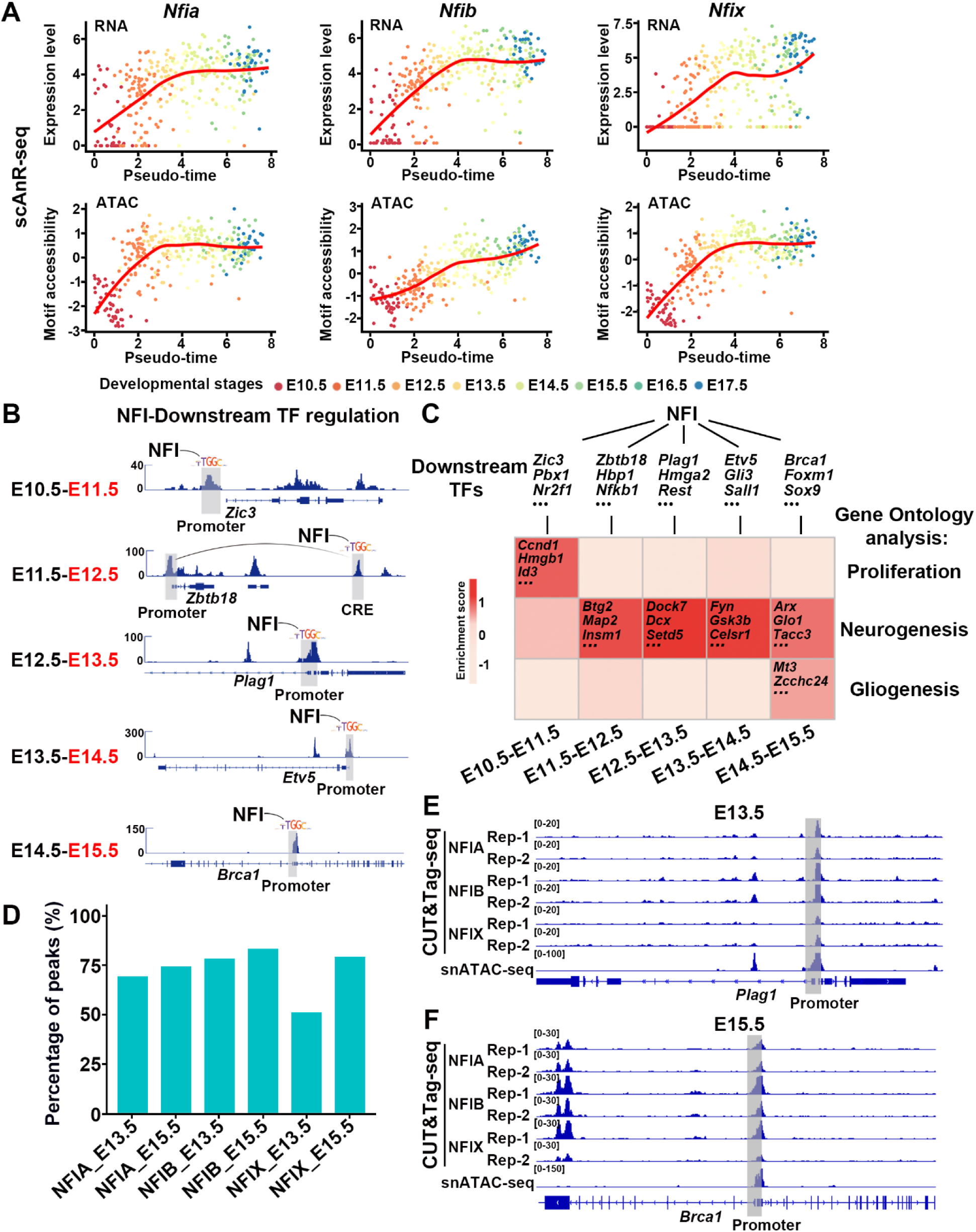
Progressive NFI activity regulates distinct temporal-specific gene networks in RGPs coupled with lineage progression. **(A)** The expression level and the regulatory target motif accessibility of NFIs (*Nfia*/NFIA, *Nfib*/NFIB, and *Nfix*/NFIX) in RGPs throughout development as assayed by scAnR-seq. Note the progressive increase of the expression level and the regulatory target motif accessibility. **(B)** Representative genome tracks showing normalized accessibility of NFI binding motifs in the promoters or cis-regulatory elements (CREs) (both shown in gray boxes) of distinct crucial downstream TFs involved in RGP development. **(C)** GO analysis of the crucial downstream TFs regulated by NFIs in RGPs throughout development. Distinct downstream TFs at different developmental stages functionally coincide with RGP lineage progression from proliferation to neurogenesis to gliogenesis. **(D)** Percentage of peaks identified jointly in CUT&Tag-seq and snATAC–seq with NFI motif analyses out of total CUT&Tag-seq peaks. The high degree of overlap demonstrates that NFIs bind extensively to regions with their motifs. **(E)** Genome browser views showing NFIA, NFIB, NFIX CUT&Tag-seq and snATAC-seq peaks in downstream TF *Plag1* promoter region at E13.5. **(F)** Genome browser views showing NFIA, NFIB, NFIX CUT&Tag-seq and snATAC-seq peaks in downstream TF *Brca1* promoter region at E15.5.

We particularly focused on TF targets of NFIs, as they often play crucial roles in regulating progenitor lineage progression and temporal development. Indeed, based on the integrated scAnR-seq dataset analysis, we found that NFIs exert global temporal control via the regulation of distinct sets of crucial downstream TFs at different developmental stages, including those at E10.5-E11.5 (e.g., *Zic3*, *Nr2f1*, and *Pbx1*), E11.5-E12.5 (e.g., *Zbtb18*, *Hbp1*, and *Nfkb1*), E12.5-E13.5 (e.g., *Plag1*, *Hmga2*, and *Rest*), E13.5-E14.5 (e.g., *Etv5*, *Gli3*, and *Sall1*), and E14.5-E15.5 (e.g., *Brca1*, *Foxm1*, and *Sox9*) **(Figures 7B and 7C)**. Moreover, the regulation by NFIs on downstream target TFs can occur either directly between NFIs and downstream TFs or indirectly through downstream TFs’ cross-regulation within their temporal network, as shown by the systematic analysis of the dynamics of accessible peaks in the promoters and/or CREs of the corresponding downstream TFs in the snATAC-seq dataset **(Figures 7B, S5D-S5I, and S6; Table S1)**. Furthermore, we performed CUT&Tag-seq experiments using NFIA, NFIB and NFIX antibodies on E13.5 and E15.5 RGPs. The NFI binding peaks identified by CUT&Tag-seq dataset exhibited high reproducibility, and showed consistent enrichment in the promoters and/or CREs of downstream TFs, aligning with the corresponding snAnR-seq peaks **(Figures 7D-7F and S5D-S5I; Table S1)**. Notably, the downstream TFs regulated by NFIs at different developmental stages exhibit distinct functional features based on gene ontology (GO) analysis of their target genes **(Figure 7C; Tables S2 and S3)**. The early targets at E10.5-E11.5 are functionally linked to progenitor proliferation, the targets at E11.5-E14.5 are more involved in neurogenesis, and the late targets at E14.5-E15.5 are functionally associated with gliogenesis. This progressive functional shift of the targets regulated by NFIs via downstream TFs aligns well with RGP lineage progression from proliferation to neurogenesis to gliogenesis. Together, these results suggest that the progressive increase in the expression of NFIs as global temporal regulators governs RGP lineage progression by regulating distinct temporal-specific gene networks including sequential sets of downstream TFs and target genes with distinct functions in controlling proliferation, neurogenesis, and gliogenesis.

### NFIA/X confer RGP temporal identity in a dose-dependent manner

The progressive expression and activity NFIs in RGPs indicate that NFIs control RGP temporal identity, which is functionally reflected in the well-defined stage-specific neural progeny output. Dividing RGPs in the WT cortex at E13.5 and E15.5 largely produce layer V and III-IV neurons, respectively, as revealed by BrdU pulse-chase labeling experiments **(Figures S7A-S7C top)**. By contrast, dividing RGPs in the *Nfia;Nfix* cDKO cortex at the same developmental stages predominantly generated layer VI and IV-V neurons, respectively **(Figures S7A-S7C bottom)**. In addition, while dividing RGPs in the E17.5 WT cortex only produced a small number of layer II neurons, those in the E17.5 *Nfia;Nfix* cDKO cortex generated a large number of layer III neurons **(Figures S7A-S7C)**. These results show that the removal of NFIA and NFIX systematically shifts the neural progeny output of RGPs toward the earlier developmental stage. That is, the temporal identity of RGPs appears younger in the absence of NFIA and NFIX.

Previous studies showed that RGPs display progressive and characteristic changes in gene transcription that reflect their developmental stage and temporal identity ^44–48^. To assess the temporal identity of RGPs, we carried out a systematic scRNA-seq analysis of WT cortices at E12.5-P7 and identified RGPs based on well-established marker gene expression **(Figures S8A and S8B)**. By employing pseudo-time analysis of RGPs to reconstruct their developmental progression, we found that RGPs obtained at different developmental stages formed an orderly trajectory from the early to late time points **(Figure 8A)**, consistent with the notion that RGPs follow a defined temporal developmental program, which is reflected in their gene expression profiles.

**Figure 8.**
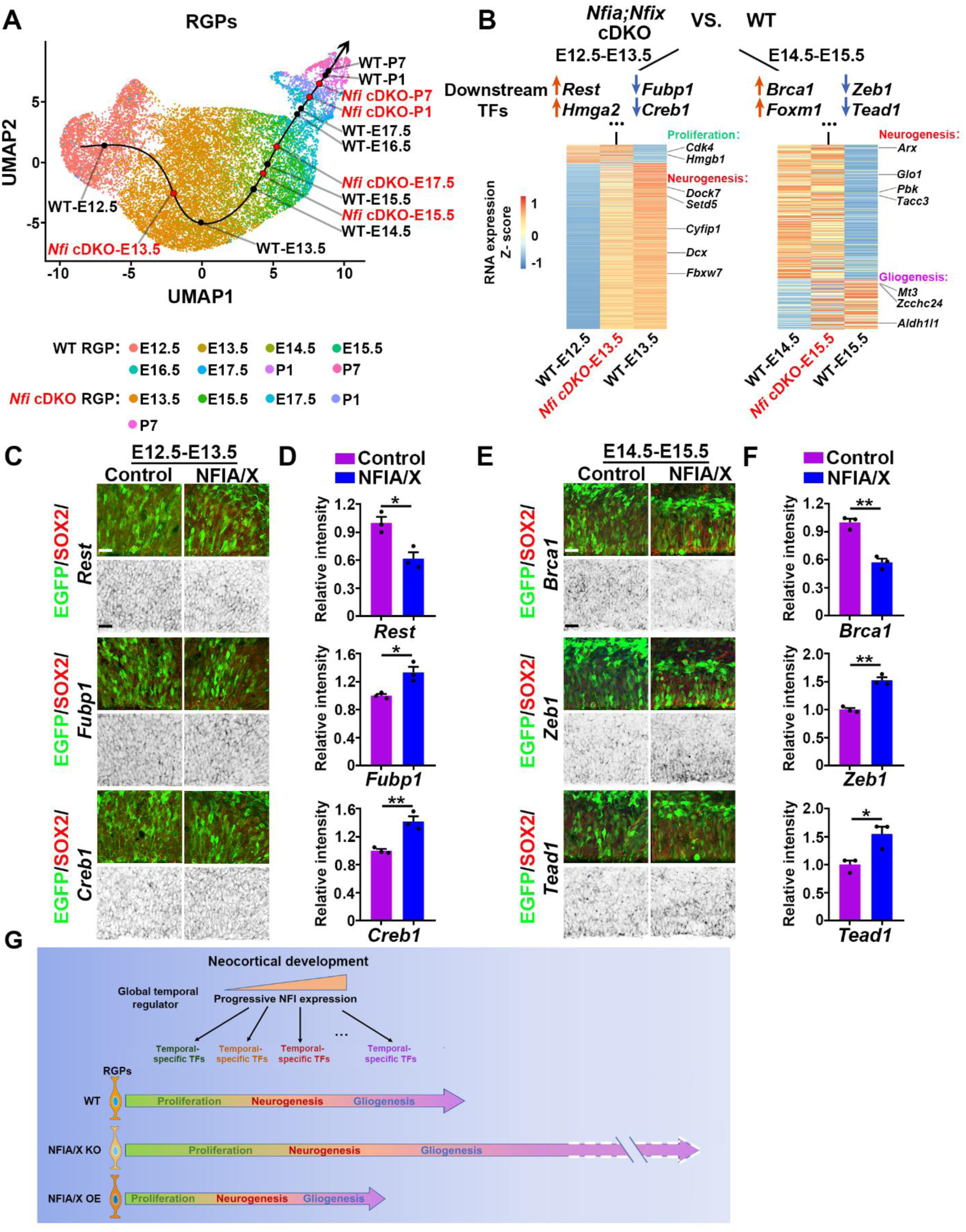
NFIA/X confer RGP temporal identity by regulating downstream TFs. **(A)** UMAP plot of RGPs from WT (E12.5, E13.5, E14.5, E15.5, E16.5, E17.5, P1 and P7) and *Nfia;Nfix* cDKO (*Nfi* cDKO) (E13.5, E15.5, E17.5, P1 and P7) cortices at different developmental stages. Each point represents an individual RGP and is labelled by distinct colors for different samples. The line represents the developmental temporal trajectory of all RGPs. Note that, compared with WT control obtained at the same developmental stage, *Nfia;Nfix* cDKO (*Nfi* cDKO) RGPs are younger in the developmental temporal trajectory. **(B)** Distinct changes in the target downstream TF expression and GO at different developmental stages upon the removal of NFIA and NFIX. GO terms associated with some target genes of downstream TFs are highlighted. Removal of NFIA and NFIX leads to an increase in the expression of genes linked to proliferation (e.g., *Cdk4 and Hmgb1*) and a concurrent decrease in the expression of genes associated with neurogenesis (e.g., *Dcx*, *Dock7*, *Setd5*, *Cyfip1*, and *Fbxw7*) at E13.5. In comparison, at E15.5 the removal of NFIA and NFIX leads to an increase in the expression of genes linked to neurogenesis (e.g., *Arx*, *Glo1*, *Pbk*, and *Tacc3*) and a concurrent decrease in the expression of genes associated with gliogenesis (e.g., *Aldh1l1, Mt3*, and *Zcchc24*). **(C)** Representative images of E13.5 cortices electroporated with EGFP control or NFIA and NFIX (NFIA/X) overexpression plasmids together with EGFP (green) at E12.5 stained for SOX2 (red). Downstream TF *Rest*, *Fubp1*, and *Creb1* mRNA (black) in situ hybridization was performed on the adjacent sections. **(D)** Quantification of the relative intensity of in situ signal in **(e)** (Control, n = 3 brains; NFIA/X, n = 3 brains). **(E)** Representative images of E15.5 cortices electroporated with EGFP control or NFIA and NFIX (NFIA/X) overexpression plasmids together with EGFP (green) at E14.5 stained for SOX2 (red). Downstream TF *Brca1*, *Zeb1*, and *Tead1* mRNA (black) in situ hybridization was performed on the adjacent sections. **(F)** Quantification of the relative intensity of in situ signal in **(G)** (Control, n = 3 brains; NFIA/X, n = 3 brains). **(G)** Summary of NFIs as global temporal regulators of RGP lineage progression rate and lifespan in cortical development. Loss or overexpression of NFIs bidirectionally tunes RGP lineage progression rate and lifespan during mammalian cortical development. Data are presented as mean ± SEM. *, P < 0.05; **, P < 0.01. Statistical analysis was performed using unpaired Student’s t-test.

To test whether NFIA and NFIX control RGP temporal identity, we performed the same scRNA-seq-based developmental progression analysis of RGPs in *Nfia;Nfix* cDKO cortices at E13.5, E15.5, E17.5, P1, and P7 **(Figures 8A and S8B)**. Similar to RGPs in WT control cortices, RGPs in *Nfia;Nfix* cDKO cortices also displayed an orderly developmental pattern from the early to late time points **(Figure 8A)**, indicating that the overall progressive development of RGPs was not disrupted by the loss of NFIA and NFIX. However, RGPs lacking NFIA and NFIX exhibited temporal identity features representing earlier developmental stages than their gestational stage **(Figure 8A)**. We quantitatively analyzed the pseudo-time of RGPs along the developmental progression in WT and *Nfia;Nfix* cDKO cortices, and found that RGPs lacking NFIA and NFIX were consistently at younger stages than WT RGPs **(Figure S8C)**. Progressive lengthening of G_1_ phase is considered a characteristic feature of RGP temporal development ^70^. Consistent with this, we found that the ratio of RGPs in G_1_ phase gradually increased in the WT cortex as time proceeded **(Figure S8D)**. Moreover, the ratios of RGPs in G_1_ phase in *Nfia;Nfix* cDKO cortices were consistently reduced compared with those in WT cortices at corresponding gestation stages, indicating a systematic shift of RGPs towards younger developmental stages in the absence of NFIA and NFIX.

To further reveal the basis of NFIA and NFIX-mediated control of RGP temporal identity, we systematically analyzed the transcriptomic features of RGPs in WT and *Nfia;Nfix* cDKO cortices across multiple developmental stages. We focused on analyzing changes in the regulation of crucial downstream TFs of NFIs identified as the key regulators of RGP lineage transition and temporal identity **(Figure 7C)**, as downstream TFs are dynamically regulated while RGPs progress through development. Compared with WT control RGPs, the magnitudes of expression changes of downstream TFs, as well as their target genes, associated with RGP lineage progression were significantly reduced in *Nfia;Nfix* cDKO RGPs **(Figures S8E and S8F; Tables S4, S5, and S6)**, indicating an attenuated regulatory dynamics that leads to a protracted RGP lineage progression. Moreover, both up- and down-regulated changes in the expression of downstream TFs were observed in *Nfia;Nfix* cDKO RGPs compared with WT control RGPs (e.g., *Rest* and *Hmga2* up-regulation, and *Fubp1* and *Creb1* down-regulation at E13.5; *Brca1* and *Foxm1* up-regulation, and *Zeb1* and *Tead1* down-regulation at E15.5) **(Figure 8B; Tables S4, S5 and S6)**, suggesting that NFIA/X can enhance or suppress the expression of downstream TFs in RGPs. Notably, the target genes of the corresponding downstream TFs in *Nfia;Nfix* cDKO RGPs displayed a systematic shift towards younger developmental stages compared with WT control RGPs **(Figures 8B)**. At E13.5, we observed an increase in the expression of target genes linked to proliferation (e.g., *Cdk4* and *Hmgb1*) and a concurrent decrease in the expression of target genes associated with neurogenesis (e.g., *Dcx*, *Dock7*, *Setd5*, *Cyfip1*, and *Fbxw7*) **(Figure 8B left; Tables S4 and S5)**. At E15.5, we observed an increase in the expression of target genes linked to neurogenesis (e.g., *Arx*, *Glo1*, *Pbk*, and *Tacc3*) and a concurrent decrease in the expression of target genes associated with gliogenesis (e.g., *Aldh1l1*, *Mt3*, and *Zcchc24*) **(Figure 8B right and Tables S4 and S6)**. Importantly, the expression levels of crucial downstream TFs linked to RGP lineage progression and temporal development exhibited opposite changes in NFIA/X overexpressing RGPs at different embryonic stages compared with those in *Nfia;Nifx* cKO RGPs **(Figures 8C-8F)**. In particular, *Rest* expression was down-regulated and *Fubp1* and *Creb1* expressions were up-regulated at E12.5-E13.5, whereas *Brca1* expression was down-regulated and *Zeb1* and *Tead1* expressions were up-regulated at E14.5-E15.5. Together, these results further support that NFIs act as global temporal regulators to regulate temporal-specific gene networks coupled with RGP lineage progression, and consequently control the temporal identity of RGPs in a dose-dependent manner.

While NFIs regulate RGP temporal development and lineage progression in both mice and humans as shown above, the lineage progression rate and lifespan of RGPs are drastically different between these two species ^38,71–73^. Given that the expression level of NFIs controls the temporal development and lineage progression of RGPs, we thereby examined the expression dynamics of NFIs in human and mouse cortical RGPs. Interestingly, human *NFIA/X* showed a much slower progressive increase in expression than mouse *Nfia/x* in RGPs during cortical development **(Figure S8G)**, consistent with a slower lineage progression rate and a longer lifespan of RGPs in humans than in mice. Since the expression of NFIs exhibits self-promoting auto-regulation **(Figures 6C, 6D, and S5C)**, we asked whether this auto-regulation displays any species difference. Indeed, we found that the strength of auto-regulation primarily depended on species-specific CRE sequences, which was weaker in humans than in mice **(Figure S8H)**. Together, these results suggest that the genetic differences in CREs affecting NFI expression and regulation contribute to diverged lineage progression pace and lifespan of RGPs across different species.

## DISCUSSION

A fundamental process in mammalian cortical development is the orderly production of a myriad of diverse cells by RGPs. This is achieved through the precise unfolding of an intricate lineage progression program, in which RGPs progressively transit from young to old age and are actively depleted towards the end while generating diverse progenies. However, how such lineage progression and the lifespan of RGPs are regulated remain not well-understood. In this study, we found that the progressive expression and activity of NFI TFs as global temporal regulators control neocortical RGP lineage progression, lifespan, and progeny output by regulating distinct temporal-specific TF targets in a dose-dependent manner **(Figure 8G)**. Reduced levels of NFI expression and activity protract RGP lineage progression and enhance progeny output, leading to an enlarged neocortex, whereas increased levels of NFI expression and activity accelerate RGP lineage progression and reduce progeny output, resulting in a diminished neocortex. Notably, these effects of NFIs on neocortical RGPs coincide well with human patient phenotypes with *NFI* misexpressions or mutations.

NFI TFs have previously been studied in the mammalian central nervous system development ^49,50,53,74,75^. They are thought to be expressed in progenitors primarily at late developmental stages and confer the capability of gliogenesis as the last step of progenitor lineage progression or other late-born progenies, such as astrocytes in the developing spinal cord ^50–52^ or bipolar interneurons and Müller glia in the developing retina ^15^. Unexpectedly, our results demonstrate that the expression and activity of NFIs progressively increase in neocortical RGPs starting from early developmental stages and regulate the entire lineage progression of RGPs from proliferation to neurogenesis to gliogenesis. Removal of NFIs not only increases the number of late-born glial cells and superficial-layer neurons, but also strongly promotes the generation of early-born deep-layer neurons, as well as progenitor proliferation. This is achieved through the regulation of dynamically expressed temporal-specific gene targets, which have distinct functions in controlling proliferation, neurogenesis, and gliogenesis. Consistent with this, overexpression of NFIs in RGPs at different developmental stages results in the generation of distinct types of neural progenies. At E12.5, more superficial versus deep layer neurons were generated. In comparison, at E15.5 and E17.5, less superficial layer neurons and more glial cells were produced. In addition, removal of NFIs leads to a protracted RGP lineage progression and a drastic increase in the production of all neural progenies. These findings highlight that NFIs function as key global temporal regulators to control the entire RGP temporal development program in the developing neocortex, and not just particular cell type(s) as suggested in the developing spinal cord or retina ^15,17^.

This difference may reflect distinct developmental contexts in which NFIs function. Alternatively, it may reflect a difference in the time window of analysis. In the developing retina, removal of NFIs also leads to sustained retinal progenitor cell proliferation and neurogenesis ^15^, consistent with our observation in the developing neocortex. However, the numbers of late-born bipolar interneurons and Müller glia in the *Nfi* mutant retina are selectively reduced compared with the WT control at P14. In comparison, we observed a dramatic increase in both early and late-born cells in the *Nfi* mutant neocortex when we extended the time window of the analysis to cover the protracted lineage progression of RGPs. It remains possible that the generation of late-born cells in the *Nfi* mutant retina is extended beyond P14. Notably, previous studies of NFIs in the developing cortex using global knockout mice that die prenatally or neonatally suggested that the removal of NFIs suppresses neurogenesis ^56,76^, in contrast to our observation that removal of NFIs enhances neurogenesis and cortical volume upon the analysis of the full lineage unfolding of RGPs using conditional mutants.

Our data point to a two-layer temporal regulatory framework of RGP lineage progression, comprising temporal-specific target TFs under the control of progressive activities of NFIs as global temporal regulators (please also see *Yuan et al., bioRxiv 2025*) ^69^. Compared with the classic temporal TF cascade well-documented in Drosophila neural progenitors with a relative short lifespan (∼20 hours) ^18^, the two-layer regulatory framework offers an additional level of global regulation on the speed or rate of progenitor lineage progression, in addition to temporal-specific TFs. This additional regulation by global temporal regulators permits further tuning of the number and diversity of progeny output. For example, evolutionarily conserved global temporal regulators, such as NFIs, regulate RGP lineage progression in both mice and humans, yet the RGP lineage progression rate and lifespan differ widely between mice and humans. In mice, RGPs progress through the temporal development program and become depleted within 2-4 weeks, whereas in humans, RGPs progress through the lineage for more than 8-9 months. This drastic difference in RGP lineage progression rate and lifespan may lie at least partly in the temporal expression dynamics of global temporal regulators like NFIs in RGPs. The faster the progressive increase in the expression of global temporal regulators, the quicker the lineage progression rate and the shorter the lifespan of progenitors should be. Indeed, we found that the progressive expression rate of NFIs in human RGPs is much slower than that in mouse RGPs. This is achieved through a weaker positive auto-regulation in expression of human NFIs than that of mouse NFIs. Together, these findings point to an interesting regulatory mechanism underlying the species-specific progenitor temporal development rate and lifespan, in conjunction with the expression of human or primate-specific genes, such as *ARHGAP11B*, *TBC1D3*, *NOTCH2NL*, and *TMEM14B*, which drives the increase in RGP lifespan and progeny output ^77–81^. It is worth noting that additional temporal regulators likely exist in addition to NFIs in regulating mammalian neural progenitor lineage progression and temporal development. Temporal gradients of RNA-binding proteins have been found in Drosophila neuroblasts to regulate crucial TF expression in postmitotic neurons and their fate specification within a lineage ^82,83^, indicating that monotonic and progressive gene expression may be a conserved mechanism for effective regulation of neural progenitor lineage and diverse neural progeny fate specification. In addition, while we showed that the expressions of crucial downstream TFs are altered upon NFI removal or overexpression, the precise mode and mechanism of regulation by NFIs remain to be elucidated. It will be interesting to explore whether both the TF activity and chromatin state regulation activity are involved. These efforts will advance the understanding of RGP lineage progression and neocortical development.

A growing number of clinical reports have revealed that haploinsufficiency (including heterozygous loss-of-function mutations or microdeletions) of *NFIs* cause macrocephaly, whereas duplications of *NFIs* lead to microcephaly ^3–11^. These observations are consistent with our findings that the removal of NFIs greatly prolongs RGP lineage progression and lifespan in mouse and human cerebral organoid models, resulting in an excessively enlarged cortex with extensive folding. Interestingly, we found that two human pathogenic variants in *NFIA* cause an increase in RGP proliferation coinciding with the macrocephaly phenotype. Conversely, overexpression of NFIs drastically accelerates RGP lineage progression, leading to precocious neural progeny output and premature loss of RGPs. Moreover, NFI overexpression can fast-forward the developmental clock of human pluripotent stem cells to switch from neurogenesis to gliogenesis in 5 days, a process that would normally take >100 days ^84^. Variable somatic overgrowth or short stature have also been observed in human patients with *NFI* deficiency or duplication, respectively, suggesting that NFIs also regulate the lineage progression and development of non-neural progenitors and tissues. Indeed, NFIs have been shown to maintain stem cell identity in skin ^68^. Together, these findings suggest that NFIs function as a key molecular control of stem and progenitor cell lineage progression and lifespan in general, raising the possibility of resetting the developmental clock of stem/progenitor cells and tuning the age of stem and progenitor cells for translational applications via NFI modulation.

## METHODS

### Animals

*Nfia^fl/fl^* (fl, floxed allele) ^63^ mice were kindly provided by Dr. Mohamed Elgazzar. *Nfib^fl/fl^* ^64^ and *Nfix^fl/fl^* ^57^ mice used in this study have been described previously. *Emx1-Cre* (stock no. 005628; The Jackson Laboratory) ^65^ and *Nex-Cre* ^67^ mice were used to delete *Nfia*, *Nfib*, and/or *Nfix*. CD-1 mice were obtained from Charles River Laboratory and Beijing Vital River Laboratory Animal Technology Co., Ltd. Genotyping was carried out using standard PCR protocols. The mice were maintained at the facilities of Tsinghua University and Memorial Sloan Kettering Cancer Center (MSKCC), and all animal procedures were approved by the Institutional Animal Care and Use Committee (IACUC). For timed pregnancies, the plug date was designated as E0.5 and the date of birth was defined as P0.

### Single Cell Library Preparation and Sequencing

For *Nfi* mutant single cell suspension preparation, E13.5, E15.5, E17.5, P1 and P7 *Nfia*;*Nfix* cDKO and littermate WT control embryos or neonatal pups were collected. In addition, E12.5, E14.5 and E16.5 WT embryos with the same background were also collected. In these experiments, two embryos or three neonatal pups per condition were used.

For collecting cortical tissues, embryonic brains were dissected and put into a dish with oxygen bubbled artificial cerebrospinal fluid (ACSF) containing: 126 mM NaCl, 3 mM KCl, 1.2 mM NaH_2_PO_4_, 1.3 mM MgCl_2_, 2.4 mM CaCl_2_, 26 mM NaHCO_3_, and 10 mM glucose (pH 7.4). Individual embryonic brains were then transferred to a new dish containing oxygen bubbled ACSF with the cortex facing up. A small area of the cortex (i.e., the dorsal medial part across the whole cortical wall from ventricle wall to pial surface) was dissected out from each hemisphere and placed into the chilled Hibernate E®/B27®/GlutaMAX™ (HEB) medium (HEB, Brainbits). For neonatal pups, cortical V-SVZ and adjacent tissues were manually dissected. The cortical tissues were dissociated following the dissociation protocol of mouse embryonic neural tissue for single-cell RNA sequencing (10X Genomics). The cell suspension was filtered using a 20 μm cell strainer (130-101-812, Miltenyi). The cell concentration and viability were measured using automated Cell Counter (cell viability >80% and cell concentration ∼700 cells/µL). Single-cell RNA-Seq was performed using Chromium™ Single Cell 3’ Library & Gel Bead Kit (10X Genomics, v3 for E12.5-E17.5 and v3.1 for P1-P7 *Nfia*;*Nfix* cDKO and littermate WT control in **Figure 8**) and Chromium™ Single Cell A Chip Kit (10X Genomics). GEM generation and library preparation were prepared according to 10X Genomics instructions.

### scRNA-seq Analysis of RGP Temporal Identity

For analyzing 10X Genomics RNA sequencing data of WT and *Nfia;Nfix* cDKO cortices, quality filtering of cell barcodes was first performed on the total counts of RNA reads and the percentage of mitochondrial genes. Cells with total counts over 1000 and mitochondrial gene percentage less than 10% were used for the subsequent analysis. Cell type identification and trajectory inference were performed using a Seurat pipeline where counts were normalized to the total reads per cell and batch variations were corrected by scaling the expression of each gene to the dataset-wide average. Datasets of different samples (i.e., different stages and genotypes) were integrated using Seurat integration method. For identifying cell types, a cell-cycle regression process was performed to limit the contribution of cell-cycle variations during clustering. Cells were classified into cell-cycle phases (G_1_, G_2_, and S) according to their RNA level of cell-cycle related genes. The top 500 variable genes were used for PCA. A UMAP method was performed to reduce dimensions to plot the top 8 principal components onto a 2-D graph. Cells were clustered based on approximate k-nearest neighbor method and cluster identities were determined by markers calculated by gene fold-change among clusters. Major cell types were identified using a common set of broad cell type marker genes. For performing pseudo-time analysis, RGP clusters were extracted, and their temporal trajectory was analyzed using Slingshot. PCA and UMAP were applied on RGP clusters. The trajectory inference was performed with setting sample birthdate and genotype as cluster identities, starting from E12.5 WT dataset and ending with P7 WT dataset. A score of pseudo-time was generated for each cell, representing the rank on the temporal trajectory.

### scAnR-seq (integrated scRNA-seq and snATAC-seq in the same individual cells)

Single cells from cortical VZs of WT CD1 mice (E10.5-E17.5) were collected for scAnR-seq as previously described ^69^. Briefly, the single cell solution was vortexed to resuspend magnetic beads and incubated on a Thermo mixer at 4℃ with shaking at 300 rpm for 30 minutes. After incubation, samples were placed on magnet for 3 minutes to perform the separation of cytoplasm and nuclei. The supernatants were carefully transferred to a new plate for scRNA-seq following Smart-seq2 protocol. The remaining nuclei were resuspended in Tn5 tagmentation mix for snATAC-seq to capture coupled chromatin accessibility for individual cells.

### *Nfi* RNA Expression and NFI Motif Accessibility Analysis in the Same Individual RGPs

Raw sequencing data from scAnR-seq were processed as previously described ^69^. A total of 361 high quality RGPs were extracted for further *Nfi* RNA expression and NFI motif accessibility analysis.

### *Nfi* RNA expression measurement

The pseudo-time axis was reconstructed based on the UMAP coordinates by Slingshot. RGPs collected from different developmental stages (E10.5-E17.5) were ordered along the pseudo-time axis sequentially. The expression patterns of *Nfi* TFs along the pseudo-time axis were fitted by ‘geom_smooth’ function in ggplot2.

### NFI motif accessibility analysis

To calculate NFI TF motif accessibility scores in individual RGPs, we adopted the method from chromVAR ^85^. Briefly, we started with the binarized cell by peak matrix and the motif annotation in each peak. The annotation for occurrence of motif was obtained by motif scan with FIMO (parameters-thresh.00001) in the Hocomoco V11 motif database ^86^. Then, we merged the fragments in all accessible peaks with the motif occurrence to form a raw cell-motif counts matrix. The cell-motif matrix was further normalized by the size factor of each cell (based on the ‘estimateSizeFactors’ function of Cicero) to correct for their differences in read depth and perform z-score for each motif. The NFI motif accessibility along pseudo-time was calculated as above TF motif accessibility scores and were ordered by pseudo-time.

### Gene Ontology (GO) Analysis

The enrichment scores for functional terms (Proliferation, Neurogenesis, and Gliogenesis) at different transition stages were calculated as follows: First, the related GO term sets of Proliferation, Neurogenesis, and Gliogenesis were collected from the GENE ONTOLOGY knowledgebase, which were correlated with each function or reported in a related study ^44^. Then, the functional enrichment for target genes of NFI via downstream TFs at each transition stage were performed by the ‘Enricher’ function of ClusterProfiler. The enrichment scores were calculated as the average -log_10_ (*P* value) of each GO term set and z-score normalized for each transition stage.

### Heatmap Analysis

To identify the differentially expressed genes between *Nfia;Nfix* cDKO and WT RGPs, we used the “FindMarkers” function of Seurat with the option (P value <0.05). The intersection of differentially expressed genes from *Nfia;Nfix* cDKO vs. WT with target genes of NFI via downstream TFs at different transition stages were plotted as a heatmap in **Figure 8B**. These differentially expressed target genes of downstream TFs were classified by the progressive changes of their expression levels between E12.5 WT vs. E13.5 WT, or E14.5 WT vs. E15.5 WT and ordered by the P values of different expression levels.

### CUT&Tag-seq

The CUT&Tag assay was performed with the Hyperactive Universal CUT&Tag Assay Kit for Illumina (Vazyme, TD903) according to the manufacturer’s instructions. Briefly, E13.5 or E15.5 cortical VZ tissues predominantly containing RGPs were dissected from wild-type CD1 mice as previously described ^87^. About 1×10^5^ cells were used for each replicate. Two replicates were performed for each antibody (rabbit anti-NFIA, HPA008884, RRID: AB_1854421, Sigma; rabbit anti-NFIB, HPA003956, RRID: AB_1854424, Sigma; mouse anti-NFIX, SAB1401263, RRID: AB_10608433, Sigma) per embryonic stage. Cells were washed and resuspended by wash buffer and then incubated with the concanavalin A-coated magnetic beads prewashed by binding buffer. Bead-bound cells were incubated with 1∼2 μg of primary antibody or nothing at 4°C overnight. After removing the primary antibody by magnetic beads, secondary antibody (goat anti-mouse or goat anti-rabbit; 1:100, Vazyme, AB206-01) was incubated at RT for 1 hour. After washing three times, the pA/G-Transposome Mix was added and incubated with bead-bound cells at RT for 1 hour to perform tagmentation. Then the DNA fragments were extracted and PCR amplification was performed using unique i5 and i7 primers from TruePrep Index Kit V2 for Illumina (Vazyme, TD202). The libraries were sequenced on NovaSeq platform with 150-bp paired-end reads. Raw fastq reads were trimmed by Cutadapt (version 1.18) to remove adapter and low-quality sequences. Fastq files were aligned to the mm10 reference genome using Bowtie2 (version 2.3.3.1; with -X 2000 option. Reads that have alignment quality < Q30, improperly paired, or aligned to ENCODE blacklisted regions (https://sites.google.com/site/anshulkundaje/projects/blacklists) were discarded. Duplicates were removed using Picard (version 2.20.4; http://broadinstitute.github.io/picard/) tools.

CUT&Tag signal for each sample were normalized to Reads Per Kilobase per Million mapped reads (RPKM). CUT&Tag peaks were called on individual samples using MACS2 peak caller ^88^ with the following parameters: -nomodel –nolambda. CUT&Tag peaks overlapped with CRE regions were calculated by the “intersect” command of Bedtools ^89^.

### Magnetic Resonance Imaging (MRI) Analysis

MRI of *ex vivo* mouse brain specimens was performed on a horizontal 7 Tesla MR scanner (Bruker Biospin) with a triple-axis gradient system. Images were acquired using a quadrature volume excitation coil (72-mm diameter) and a receive-only 4-channel phased array cryogenic coil. The specimens were prepared as described in our *Bio-protocol* paper ^90^. High-resolution diffusion MRI data were acquired using a modified three-dimensional (3D) diffusion-weighted gradient and spin echo (DW-GRASE) sequence ^91^ with the imaging parameters described previously ^90^.From the diffusion MRI data, diffusion tensors were calculated using the log-linear fitting method implemented in DTIStudio (http://www.mristudio.org) at each pixel, and a fractional anisotropy (FA) map was calculated from diffusion tensor data. Due to the large morphological differences between the WT and *Nfia;Nfix* cDKO brains, manual segmentation of the whole brain and neocortex was performed on the average diffusion weighted image and FA map, which provided tissue contrasts to delineate basic brain compartments and gray matter and white matter boundaries. Structural volumes were calculated from the segmentation results as the number of voxels in each structure and voxel volume.

### Tissue Preparation, Immunohistochemistry, and Imaging

For embryonic tissue preparation, timed pregnant females were anesthetized and embryos were removed and perfused with ice-cold PBS (pH 7.4), followed by 4% PFA. Brains were post-fixed with 4% PFA for 6 hours, cryo-protected, and sectioned at 20 μm for immunohistochemistry. Postnatal animals were similarly processed and cryo-sectioned at 40 μm. Sections were blocked in 10% serum with 0.5% Triton-X in PBS and incubated with the primary antibody at 4°C overnight. Organoids at D30 or D60 were isolated and fixed in 4% PFA overnight on a rocker, and then washed with PBS and submerged in a 30% sucrose solution until the organoids sank to the bottom of the tube. Organoids were then embedded and frozen in OCT compound (Sakura) and sectioned on a cryostat at 12 μm. The following primary antibodies were used: mouse anti-PAX6 (SC-81649; RRID: AB_1127044; 1:100; Santa Cruz), rabbit anti-PAX6 (PD022; RRID: AB_1520876; 1:500; MBL), rabbit anti-CUX1 (SC-13024; RRID: AB_2261231; 1:200; Santa Cruz), mouse anti-Ki67 (556003; RRID: AB_396287; 1:1000; BD), rat anti-BrdU (AB6326; RRID: AB_305426; 1:500; Abcam), rat anti-CTIP2 (18465; RRID: AB_2064130; 1:500; Abcam), rat anti-TBR2 (14-4875-82; RRID: AB_11042577; 1:100; eBioscience), mouse anti-TBR2 (14-4877-82; RRID: AB_2572882; 1:100; eBioscience), rabbit anti-BLBP (AB32423; RRID: AB_880078; 1:500; Abcam), mouse anti-TUJ1 (MMS-435P; RRID: AB_2313773; 1:500; Covance), rabbit anti-OLIG2 (AB9610; RRID: AB_570666; 1:500; Millipore), rabbit anti-S100 (Z0311; RRID: AB_10013383; 1:1000; Dako), chicken anti-GFP (GFP-1020; RRID: AB_10000240; 1:2000;Aves), rabbit anti-NFIA (HPA008884; RRID: AB_1854421; 1:300; Sigma), rabbit anti-NFIB (HPA003956; RRID: AB_1854424; 1:300; Sigma), rabbit anti-NFIX (NBP2-15038; RRID: AB_2891313; 1:300; Novus), mouse anti-NFIX (SAB1401263; RRID: AB_10608433;1:150; Sigma), and mouse anti-RC2 (RC2; RRID: AB_531887; 1:20; DSHB), mouse anti-SOX2 (66411-1-Ig; RRID:AB_2881783; 1:500; Proteintech), goat anti-SOX2 (AF2018; RRID: AB_355110; 1:500; R&D System), rat anti-SOX2 (14-9811-82; RRID: AB_11219471; 1:500; Invitrogen) and, rabbit anti-PHH3 (9701; RRID: AB_331535; 1:1000; Cell Signaling). The following secondary antibodies were used: donkey anti-chicken IgY (H + L) 488 (703-546-155; RRID: AB_2340376; 1:1000; Jackson ImmunoResearch), donkey anti-mouse IgG (H + L) 555 (A-31570; RRID: AB_2536180; 1:1,000; Thermo Fisher Scientific), donkey anti-mouse IgG (H+L) 488 (A-21202; RRID: AB_141607; 1:1000; Thermo Fisher Scientific), donkey anti-mouse IgM 488 (715-545-140; RRID: AB_2340845; 1:1000; Jackson ImmunoResearch), donkey anti-mouse IgG (H+L) 647 (A-31571; RRID: AB_162542; 1:1000; Thermo Fisher Scientific), donkey anti-goat IgG (H+L) 488 (A-11055; RRID: AB_2534102; 1:1000; Thermo Fisher Scientific), donkey anti-rat IgG (H+L) Cy3 (713-165-150; RRID: AB_ 2340666; 1:1000; Thermo Fisher Scientific), donkey anti-rat IgG (H+L) 488 (A-21208; RRID: AB_2535794: 1:1000; Thermo Fisher Scientific), donkey anti-rabbit IgG (H + L) 555 (A-21432; RRID: AB_141788; 1:1000; Thermo Fisher Scientific), donkey anti-rabbit IgG (H + L) 488 (A21206; RRID: AB_ 2535792; 1:1000; Thermo Fisher Scientific), and donkey anti-rabbit IgG (H + L) 647 (A-31573; RRID: AB_2536183; 1:1000; Thermo Fisher Scientific). Nuclei were stained with DAPI (D1306; RRID: AB_2629482; 1:1000; Thermo Fisher Scientific). EdU staining was performed using Click-IT EdU Alexa Fluor 647 Imaging Kit (Thermo Fisher Scientific). BrdU staining was performed as described previously ^92^. Coronal sections were mounted on glass slides, imaged with a FV1200 or FV3000 confocal microscope (Olympus), NanoZoomer 2.0 HT slide scanner (Hamamatsu Photonics) and Axio Scan.Z1 slide scanner (Zeiss), and analyzed with Photoshop (Adobe Systems) and ImageJ.

### Luciferase reporter assay

The DNA fragments of mouse *Nfi*/human *NFI* promoter, mouse *Nfi* or *Zbtb18* putative enhancer (cis regulatory element, CRE) were amplified by PCR and subsequently cloned into pGL4.10 or pGL4.23 firefly luciferase vector (Promega) upstream of Luc2 gene, respectively.

The primers used for amplifying the mouse *Nfi* promoter in **Figures 6C, 6D and S5C** were:

*Nfia* Promoter_F: ACCTCTCCCGACGATACTATTCCCAT

*Nfia* Promoter_R: GCTCTTGCCGCTCCACGCGCTGAACTTTA

*Nfib* Promoter_F: TAAACGAAGAGTGGCAGAGAGAAG

*Nfib* Promoter_R: AAACCCAGTCCTCCTTAAATAGCC

*Nfix* Promoter_F: GAGAGAGTTTTTACAGAACCTGCC

*Nfix* Promoter_R: GGCAGTACGGGGAGTACAT

The primers used for amplifying the mouse *Nfi* and *Zbtb18* CRE in **Figures 6C, 6D, S5C and S5I** were:

*Nfia* CRE_F: CACATTGTATTGGTTAGCATGGAC

*Nfia* CRE_R: CACTAGATCAGGAAGAGGAAACG

*Nfib* CRE_F: CATTTATTAGCAAGCAACTGTGCG

*Nfib* CRE_R: ACAAAACACTGGAAAGGGAGAGTT

*Nfix* CRE_F: TAGGTTTTTGTGTGGAACCTGGAA

*Nfix* CRE_R: GGACCCTCATTTTGTTGTGATGAT

*Zbtb18* CRE_F: CGGAACAGTTGGTGAGCTAA

*Zbtb18* CRE_R: TGAAGGTGTGCGCTGAGTGA

The primers used for amplifying the mouse *Nfia* CRE (m*Nfia* CRE) and human *NFIA* CRE (h*NFIA* CRE) in **Figure S8H** were:

m*Nfia* CRE _F: ACCTCTCCCGACGATACTATTCCCAT

m*Nfia* CRE _R: GCTCTTGCCGCTCCACGCGCTGAACTTTA

h*NFIA* CRE _F: ACCTCTGCCGACGAATCTATTCC

h*NFIA* CRE _R: GCTATTGCCGCTCCATGAGC

Human NFIA cDNA was amplified by PCR and cloned into pCAG-IRES-EGFP vector, using the following primers:

NFIA (human)_F: ATGTATTCTCCGCTCTGTCTCAC

NFIA (human)_R: TTATCCCAGGTACCAGGACTGT

For luciferase reporter assay, HEK293T cells were seeded into 24-well plates and co-transfected with the luciferase reporter vectors and NFI expression plasmids. Around 48 h after transfection, cells were harvested for luciferase assay using the Dual-Luciferase Reporter Assay System (Promega, E1910) according to the manufacturer’s instructions. For **Figure 6C, 6D and S5C**, the amounts of plasmids for each transfection were: 40 ng pGL4.73 (Promega, Renilla luciferase vector), 240 ng pCAG-IRES-EGFP control or mouse NFIA/NFIB/NFIX expression vectors (pCIG-NFIA/pCAGG-NFIB2/pCIG-NFIX), 240 ng pGL4.23-empty (pGL4.10-empty) (Promega) or pGL4.23-mouse *Nfi* CRE (pGL4.10-mouse *Nfi* Promoter). For **Figure S5I**, the amounts of plasmids for each transfection were: 40 ng pGL4.73 (Promega, Renilla luciferase vector), 240 ng pCAG-IRES-EGFP control or 80 ng each of mouse NFIA, NFIB and NFIX expression vectors (pCIG-NFIA, pCAGG-NFIB2 and pCIG-NFIX), 240 ng pGL4.23-empty (Promega) or pGL4.23-mouse *Zbtb18* CRE. For **Figure S8H**, the amounts of plasmids for each transfection were: 40 ng pGL4.73 (Promega, Renilla luciferase vector), 240 ng pCAG-IRES-EGFP control or mouse/human NFIA expression vectors (pCIG-mouse NFIA/pCIG-human NFIA), 240 ng pGL4.10-empty (Promega) or pGL4.10-mouse *Nfia* CRE/pGL4.10-human *NFIA* CRE. Firefly luciferase activity was measured by Varioskan Flash spectral scanning multimode reader (Thermo Scientific) and normalized to Renilla luciferase activity in all conditions.

### Pair-Cell Analysis

Pair-cell analysis was performed as previously described ^93^. In brief, E13.5 cortices were dissected and enzymatically dissociated using 2 mg/mL papain solution (PAP/HE, Brainbits) at 37°C for 20 minutes. Then the tissue was pipetted to generate single cell suspension. Cells were plated on poly-D-lysine (P9155, Sigma Aldrich) coated 96-well plates at low density and cultures were maintained at 37°C with constant 5% CO_2_ supply. Around two hours later retroviruses carrying EGFP were added to cultures, and the cultures were fixed 24 hours later for immunostaining and further cell fate analysis of EGFP^+^ cell pairs.

### Cell Cycle Exit Analysis and BrdU Labeling

To label proliferating cells, pregnant females were intraperitoneally injected with EdU (10 µg per gram body weight). At 24 hours after the injection, embryonic brains were collected, sectioned, and stained with the antibodies to Ki67. EdU staining was performed according to the manufacturer’s protocol (Thermo Fisher Scientific, C10340). The cell cycle exit index was analyzed as the fraction of EdU^+^;Ki67^-^ cells among all EdU^+^ cells.

Mice (e.g., P60) were administered with a single intraperitoneal injection of BrdU (100 µg per gram body weight) and were sacrificed 2 hours after BrdU injection. For BrdU birthdating of neurogenesis, timed pregnant female mice (e.g., E13.5, E15.5, or E17.5) or P1 mice were administered with a single intraperitoneal injection of BrdU (50 µg per gram body weight) and were sacrificed at P21. For BrdU birthdating of gliogenesis, P60 mice were administered twice daily with intraperitoneal injections of BrdU (100 µg per gram body weight) for 3 days (P60-P62) and were sacrificed 5 weeks later (P97).

### In Utero Intraventricular Injection and Electroporation

Mouse *Nfia* or *Nfix* cDNA was inserted into a pCAG-IRES-EGFP vector for overexpression. For in utero electroporation, 1–1.5 μL of plasmids (3 μg/μL) mixed with Fast Green were injected into the embryonic cerebral ventricle through a beveled, calibrated glass micropipette. Electroporation was carried out using an electroporator (ECM830, BTX Harvard Apparatus) (5 pulses, 35 Volts for E12.5; 38 Volts for E14.5; 40 Volts for E15.5; 45 Volts for E17.5; 50 msec duration, 950 msec interval). After electroporation, the uterus was placed back in the abdominal cavity and the wound was surgically sutured. After surgery, the animal was placed in a recovery incubator under close monitoring until it fully recovered.

### RNA in situ hybridization

RNA in situ hybridization experiments were performed as previously performed using digoxigenin riboprobes on 20 μm frozen sections ^92^. Riboprobes were amplified by PCR from embryonic mouse cDNA using the following primers:

*Fubp1*_F: GGATAGCACAGATAACAGGACCTC

*Fubp1*_R: TTGTCCGTTCGTTTGGGTTGTAG

*Creb1*_F: CTGATTCCCAAAAACGAAGGGAAA

*Creb1*_R: CATTTTCCTCATTTCCCCCAACAA

*Rest*_F: GCAAATCAAAAATCGGTACCAACG

*Rest*_R: TTTTGTTCAGAGGATGAAACACGG

*Zeb1*_F: TCCGATGATGAAGACAAACTCCAT

*Zeb1*_R: GGGTTGAACAGTTGATTCCTGAAG

*Tead1*_F: AGACTCGTACAACAAACACCTCTT

*Tead1*_R: GCATCACAACTTCACAGCTACAAT

*Brca1*_F: ATAATCCCAGATTCAGAGGCATCC

*Brca1*_R: AACTATCCACTTTCCTCCTGCAAT

### Generation of *NFI* KO and *NFIA* patient variant mutation hESCs and Cerebral Organoid Generation

H1 ES cells were tested for karyotype and mycoplasma before targeting. To generate the *NFIA;NFIX* DKO hESCs, guide RNA (gRNA) targeting human *NFIA* (ATTCCGCTGGAAAGTACTGA) and *NFIX* (CCCCATCAGTACTT TCCAGG) were nucleofected alongside Cas9 RNPs. Clones were picked and expanded and indels were confirmed by PCR of the genomic DNA.

To generate the *NFIA* patient variant mutation hESCs, CRISPR/Cas9 mediated homology-directed repair was used to generate patient specific mutations in H1 ES cells. Patient 1 ^19^ has one c.220 C>T (p.Arg74Ter) mutation in exon 2 in *NFIA*. A gRNA (5’- GAAGTGGGCATCTCGACTTC-3’) was used to generate double strand break, and a donor DNA template (5’-ACATGAAAAGCGTATGTCAAAAGAAGAAGAGAGAGCCGTGAAGGATG AATTGCTAAGTGAAAAACCAGAGGTCAAGCAGAAGTGGGCATCTTGATTGCTGGCA AAGTTGCGGAAAGATATCCGACCCGAA-3’) was used to guide homology-directed repair to introduce the patient mutation and a synonymous mutation that prevent repeated binding of Cas9/gRNA. Single clones were picked, expanded and validated by Sanger Sequencing (Forward Primer: 5’-CATCCTTTCATCGAAGCACTTCTGC-3’; Reverse Primer: 5’- ACTTTATCTGCC TGGCGGAGG-3’).

Patient 2 ^20^ was reported to have a pathogenic variant c.373A>G (p.Lys125Glu) also in exon 2 in NFIA. A separate gRNA (5’- TCCGCCAGGCAGATAAAGTC-3’) and donor DNA template (5’-CAGTTACAGGGAAAAAACCTCCATGTTGTGTTCTTTCCAACCCAGA CCAGAAAGGCAAGATGCGAAGAATTGACTGCCTCCGCCAGGCAGATGAGGTTTGGAGGTTGGACCTTGTTATGGTGATTTTGTT-3’) was used to generate this mutation, also validated by Sanger Sequencing (Forward Primer: 5’- GGTCAAGCAGAAGTGGGCATC-3’; Reverse Primer: 5’- CTGCTGCATGCACAAAGTATGCC-3’). Cerebral organoids were generated as previously described ^58,59^ with some modifications. Briefly, hESCs were maintained in Essential 8 (E8) media (passages between 35-50). hESCs were dissociated by Accutase and replated to achieve ∼2,000 cells per spheroid in either 96 well V-bottom dishes (S-bio) or Aggrewell 400 wells in E8 media with the addition of 10 µM Y27632 (day 0, D0, of differentiation). Embryoid body formation media was added to the spheroids for a total of 4 days after (D5) and switched to neural induction media for an additional 5 days (D10). Spheroids were then embedded in 100% Matrigel either in the V-bottom dish directly, or when Aggrewells were used, a Matrigel sheet-style embedding technique was used. The embedded spheroids were then transferred to ultra-low attachment dishes and cultured on a shaker upon harvesting at D30 and D60 of the differentiation.

### NFIA and NFIX Overexpression hESC-derived Cerebral Organoids

*NFIA* and *NFIX* cDNAs were cloned from human neural stem cells and inserted into the FUW lentiviral tetracycline inducible system ^84^. Viruses were generated by transfecting 293T cells using the PEI transfection reagent. H9 ES cells expressing a stable rtTA transgene were infected by the viruses. Cerebral organoids were generated as described above and on D10 of the differentiation, cells were either treated with 2 µg/mL doxycycline or medium control.

### qRT-PCR Expression Analysis of Regional Markers in Cortical Organoids

Cortical organoids were harvested after 20 days of culture (D20) and placed in TriZol. To ensure that RNA was extracted from the entire organoid, each sample was passed through a 22-gauge syringe to break up the tissue. RNA was extracted using an RNEasy mini prep kit (Qiagen) and 1 µg of cDNA was generated using SuperscriptVILO (Invitrogen) per the manufacturer’s instructions. The resulting cDNA was used for qRT-PCR reactions using the SsoFast EvaGreen mix (Biorad). The primer sequences for the qRT-PCR are as follows:

*FOXG1*_F: CAACGGCATCTACGAGTTCA

*FOXG1*_R: TGTTGAGGGACAGATTGTGG

*OTX2*_F: AGAGGAGGTGGCACTGAAAA

*OTX2*_R: GCTGTTGTTGCTGTTGTTGG

*HOXB*4_F: CTGGATGCGCAAAGTTCAC

*HOXB4*_R: CCTTCTCCAGCTCCAAGACC

*PAX6*-F: GTCCATCTTTGCTTGGGAAA

*PAX6*-R: TAGCCAGGTTGCGAAGAACT

*GSX2*_F: AGATTCCACTGCCTC

*GSX2*_R: CAGGAGTTGCGTGCTAGTGA

*NKX2.1*_F: ATGTGGTCCGGAGGCAGT

*NKX2.1*_R: TGCTTTGGACTCATCGACAT

*TBR1*_F: CACAACTGAAAATAGATCACAACC

*TBR1*_R: GTCAGGCGGTCCATGTCAC

*CTIP2*_F: GACTCAGGGTGAGGGTCAGA

*CTIP2*_R: GGCTGCTTGCATGTTGTG

*TBR2*_F: CGGCCTCTGTGGCTCAAA

*TBR2*_R: AAGGAAACATGCGCCTGC

### Quantification and Statistical Analysis

Age-matched WT littermates were used as the control in all experiments. For cell number quantification, all cells positive for the corresponding markers were counted in columnar areas with indicated widths provided in the figure legends from the lateral ventricle to the pial surface of the cortex per brain section. The number of times each experiment was repeated independently with similar results is provided in the figure legends. Both male and female mice were used in the experiments. All statistical tests were performed with Prism (GraphPad). Data were presented as mean ± SEM (standard error of the mean) or mean ± SD (standard deviation of the mean) and statistical differences were determined using One-way ANOVA, Two-way ANOVA, unpaired Student’s t-test, Chi-square test, or pairwise Wilcoxon rank sum test. Statistical significance was set as p < 0.05.

## ACKNOWLEDGEMENTS

We thank the members of the Shi and Joyner laboratories, and Dr. Kun Huang (Indiana University, USA) for valuable discussion and input at the early stage of the study. Dr. N.S. Bayin for valuable input and discussion on *Nfi* gene function. Dr. Z. Yang and Dr. Y. Xie for sharing luciferase plasmids and antibodies. This work was supported by STI2030-Major Projects (2021ZD0202300) (S.-H.S.), National Natural Science Foundation of China (32021002) (S.-H.S.), Beijing Outstanding Young Scientist Program (BJJWZYJH01201910003012) (S.-H.S.), Beijing Municipal Science & Technology Commission (Z211100003321001 and Z221100003422011) (S.-H.S.), Chinese Institute for Brain Research (Beijing) (S.-H.S.), New Cornerstone Investigator Program (S.-H.S.), Tsinghua University Initiative Scientific Research Program (S.-H.S.), the NIH (R01DA024681 to A.L.J. and P30CA008748 to Memorial Sloan Kettering Cancer Center), the Tri-SCI (2019-022 to A.L.J. and S.-H.S.), the National Key R&D Program of China (2024YFA09196, 2024YFA09173, 2019YFA0904400, 2019YFA0906700) (Y.L.), Beijing Natural Science Foundation (Z210010) (Y.L.), the National Natural Science Foundation of China (32171448) (Y.L.), Tsinghua University Initiative Scientific Research Program (2024Z11DSZ001, 2022Z11QYJ032, 2021Z11JCQ020) (Y.L.), and NIH Pioneer award (DP1OD031273) (L.J.R.).

## AUTHOR CONTRIBUTIONS

Q.Z. and S.-H.S. conceived the project; Q.Z. performed the majority of the experiments and data analysis with extensive support from G.Y., J.Y., X.Y., Z.Z., E.A., X.C., C.H.L., H.D., L.Z., K.A., X.Z., X.L., X.T., S.H., Y.C., J.M., R.M.G., L.J.R., J.Z., A.L.J., J.T., and Y.L.; In particular, Q.Z. generated mutant mice and other samples, and performed experiments as well as data analysis with support from Z.L., A.K., X.Z., X.L., S.H., Y.C., J.M., and A.L.J.; R.M.G. and L.J.R. provided mouse samples; L.J.R. shared unpublished data and expertise on *Nfi* gene function in RGC development; C.H.L., and J.Z. performed MRI experiment and data analysis. G.Y., J.Y., X.Y., Z.Z., X.T., and Y.L. performed bioinformatic analysis related to scAnR-seq, CUT&Tag-seq and 10x scRNA-seq; E.A., X.C., and J.T. performed human cortical organoid-related experiment and data analysis. H.D., performed in situ hybridization experiment. This study was supervised by A.L.J., J.T., Y.L., and S.-H.S. Q.Z. and S.-H.S. wrote the paper with inputs from all authors.

---

**Figure S1.**
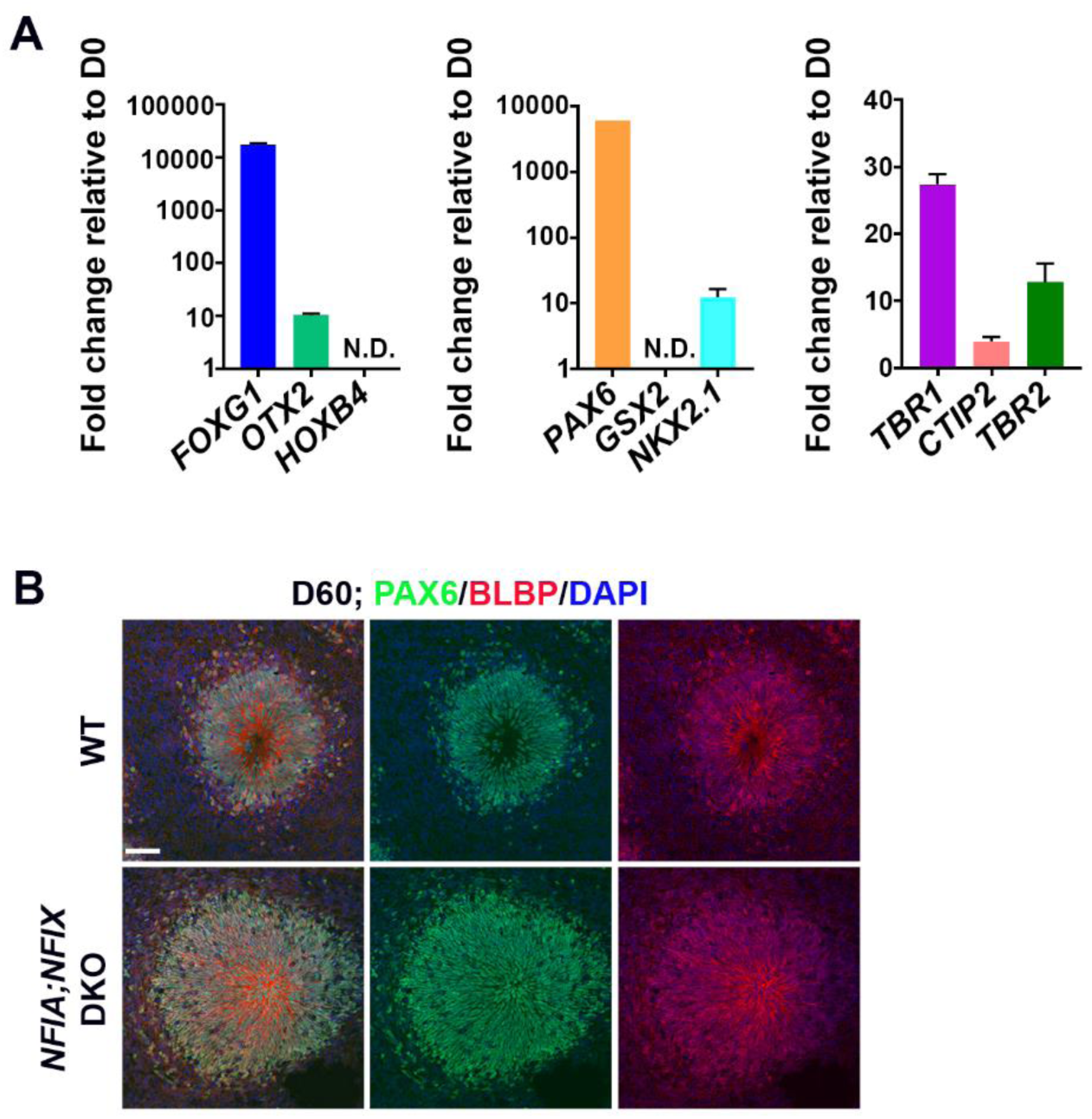
Manipulation of NFI expression bidirectionally tunes human RGP lineage progression, related to. **Figure 1 and 2**. **(A)** Quantitative PCR analysis of the fold changes of brain regional markers in human cerebral organoids at day (D) 20 compared with D0 (n = 3 biological replicates, 10 organoids pooled). **(B)** Representative images of D60 WT and *NFIA;NFIX* DKO human cerebral organoid sections stained for PAX6 (green) and BLBP (red), and with DAPI (blue). Scale bar, 50 µm.

**Figure S2.**
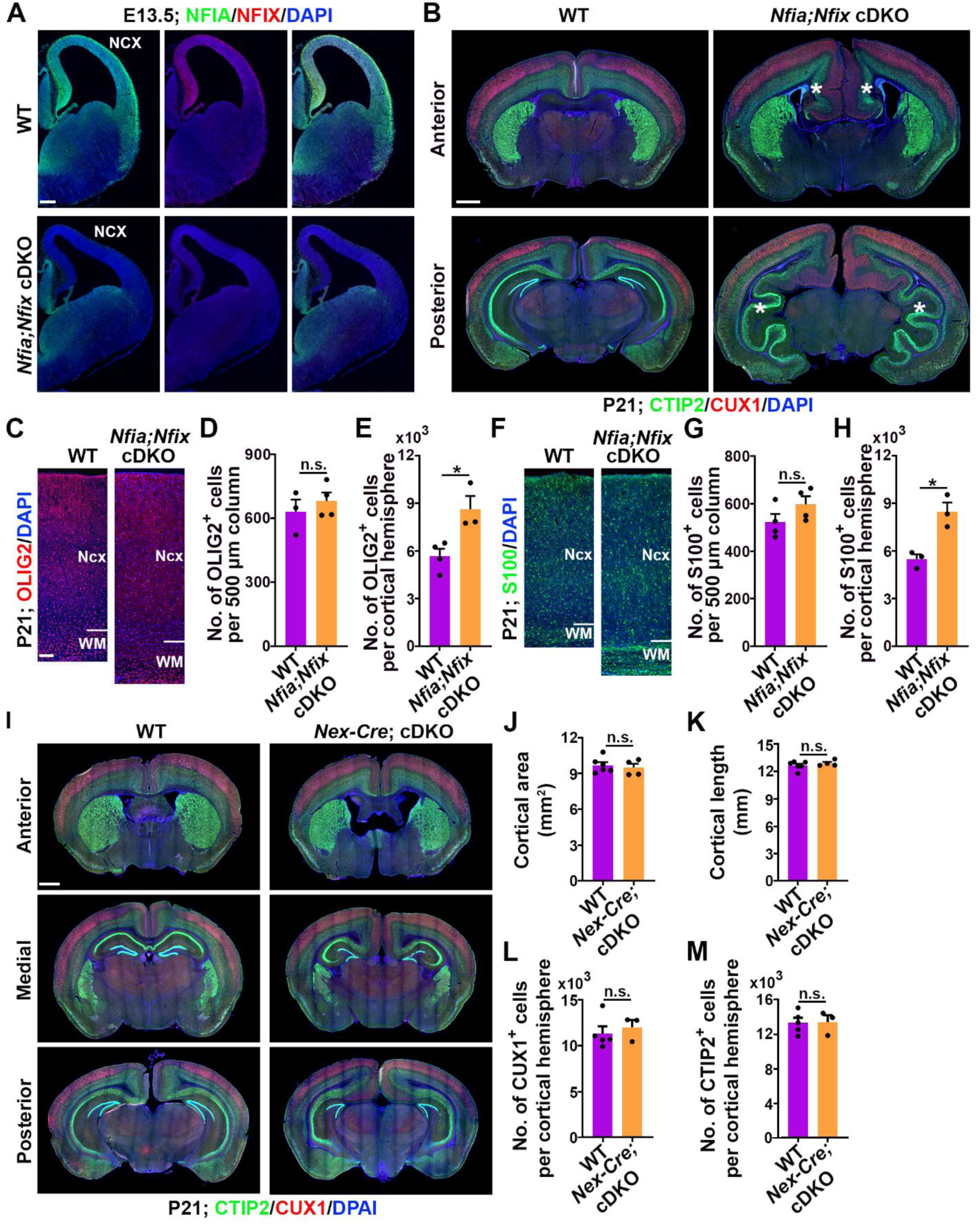
The removal of NFIA and NFIX in RGPs causes extensive cortical expansion, related to
Figure 3. **(A)** Representative images of E13.5 wild-type (WT) and *Nfia;Nfix* cDKO cortices stained for NFIA (green) and NFIX (red), and with DAPI (blue). Note the loss of NFIA and NFIX expression in the dorsal cortex in the *Nfia;Nfix* cDKO brain. Ncx, neocortex. Scale bar, 200 µm. **(B)** Representative images of P21 WT and *Nfia;Nfix* cDKO brain sections stained for CTIP2 (green) and CUX1 (red), and with DAPI (blue). White asterisks indicate the dramatic folding in the medial cortex and the hippocampus. Note the agenesis of corpus callosum in *Nfia;Nfix* cDKO brain. Scale bar, 1 mm. **(C)** Representative images of P21 WT and *Nfia;Nfix* cDKO brain sections stained for OLIG2 (red), an oligodendrocyte marker, and with DAPI (blue). Ncx, neocortex; WM, white matter. Scale bar, 100 µm. **(D)** Quantification of the number of OLIG2^+^ oligodendrocytes per 500 μm width column in the dorsal neocortex (WT, n = 3 brains; *Nfia;Nfix* cDKO, n = 4 brains). **(E)** Quantification of the total number of OLIG2^+^ oligodendrocytes per cortical hemisphere (WT, n = 4 brains; *Nfia;Nfix* cDKO, n = 3 brains). **(F)** Representative images of P21 WT and *Nfia;Nfix* cDKO brain sections stained for S100 (green), an astrocyte marker, and with DAPI (blue). Ncx, neocortex; WM, white matter. Scale bar, 100 µm. **(G)** Quantification of the number of S100^+^ astrocytes per 500 μm width column in the dorsal neocortex (WT, n = 4 brains; *Nfia;Nfix* cDKO, n = 4 brains). **(H)** Quantification of the total number of S100^+^ astrocytes per cortical hemisphere (WT, n = 3 brains; *Nfia;Nfix* cDKO, n = 3 brains). **(I)** Representative images of P21 WT and *Nex-Cre;Nfia;Nfix* cDKO brain sections stained for CTIP2 (green) and CUX1 (red), and with DAPI (blue). Scale bar, 1 mm. **(J and K)** Quantification of the cortical area **(J)** and length **(K)** (WT, n = 6 brains; *Nex-Cre; Nfia;Nfix* cDKO, n = 4 brains). **(L and M)** Quantification of the total number of CUX1^+^ **(L)** and CTIP2^+^ **(M)** cortical neurons per hemisphere in the WT and *Nex-Cre;Nfia;Nfix* cDKO brains at P21 (WT, n = 5 brains; *Nex-Cre;Nfia;Nfix* cDKO, n = 3 brains). Data are presented as mean ± SEM. *, P < 0.05; n.s., not significant. Statistical analysis was performed using unpaired Student’s t-test.

**Figure S3.**
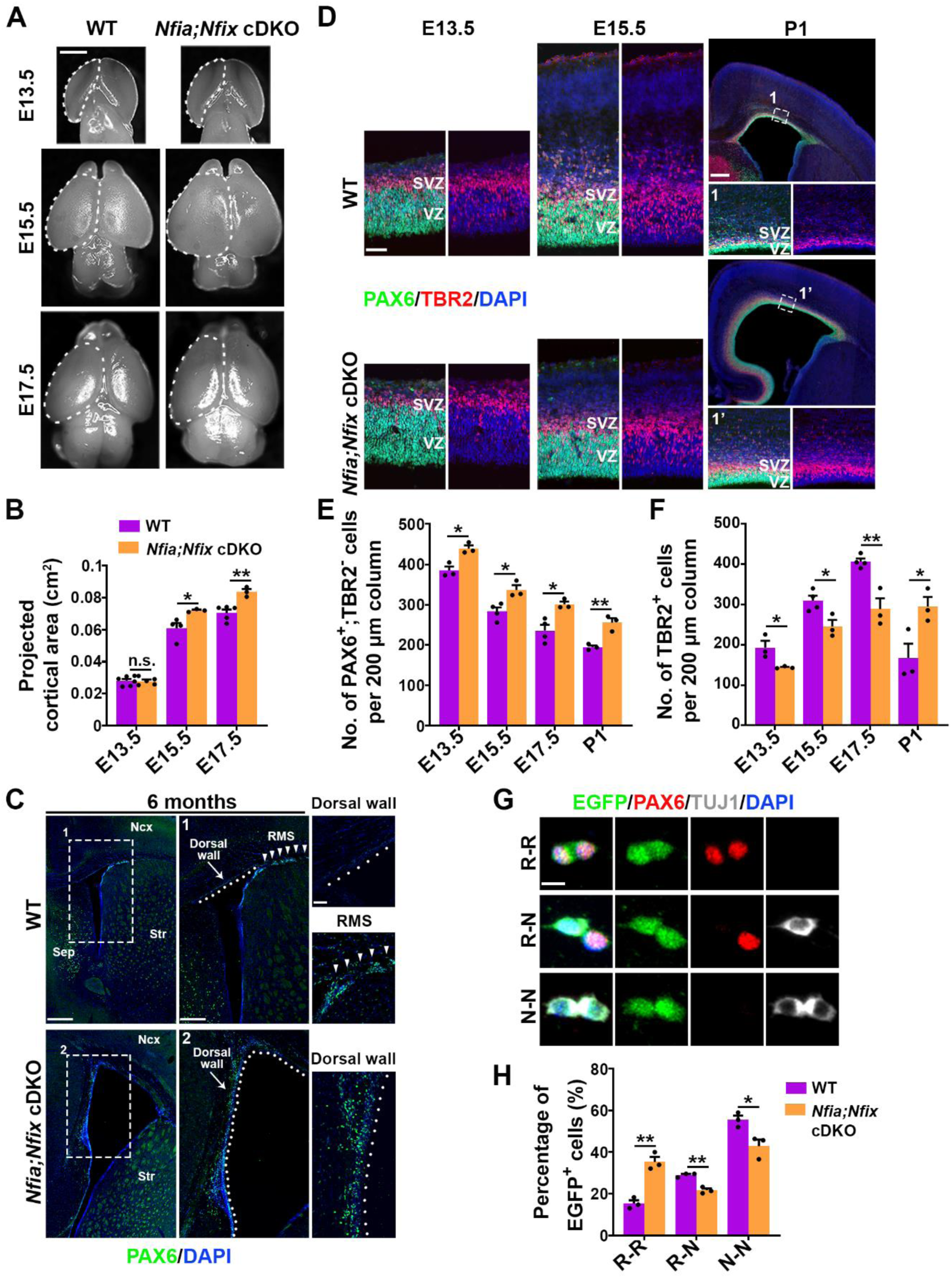
The removal of NFIA and NFIX in RGPs leads to a dramatic increase in RGPs and a concomitant decrease in IPs at the embryonic stage, related to Figure 4. **(A)** Whole mount images of E13.5, E15.5, and E17.5 WT and *Nfia;Nfix* cDKO brains. Scale bar, 1 mm. **(B)** Quantification of the projected cortical area (E13.5: WT, n = 6 brains; *Nfia;Nfix* cDKO, n = 5 brains; E15.5: WT, n = 4 brains; *Nfia;Nfix* cDKO, n = 3 brains; E17.5: WT, n = 5 brains; *Nfia;Nfix* cDKO, n = 3 brains). **(C)** Representative images of 6-month-old WT and *Nfia*;*Nfix* cDKO cortices stained for PAX6 (green), an RGP marker, and with DAPI (blue). The dashed lines indicate the boundaries of the dorsal wall of the lateral ventricle. Note the extensive presence of PAX6^+^ RGPs in the dorsal wall of the lateral ventricle in the *Nfia*;*Nfix* cDKO cortex, compared with the WT control. Ncx, neocortex; Str, striatum; Sep, septum; RMS, rostral migratory stream. Scale bars, 500 µm (left), 250 µm (middle), 50 µm (right). **(D)** Representative images of E13.5, E15.5, and P1 WT and *Nfia;Nfix* cDKO cortices stained for PAX6 (green) and TBR2 (red), and with DAPI (blue). Scale bars, 500 µm (left) and 250 µm (right). **(E and F)** Quantifications of the number of PAX6^+^;TBR2^-^ (RGP) **(E)** and TBR2^+^ (IP) **(F)** cells per 200 μm width area in the medial cortex (E13.5: WT, n = 4 brains; *Nfia;Nfix* cDKO, n = 4 brains; E15.5: WT, n = 5 brains; *Nfia;Nfix* cDKO, n = 3 brains; E17.5: WT, n = 4 brains; *Nfia;Nfix* cDKO, n = 3 brains; P1: WT, n = 3 brains; *Nfia;Nfix* cDKO, n = 3 brains). **(G)** Representative images of three different types of daughter-cell pairs originating from individual EGFP-expressing WT or *Nfia;Nfix* cDKO RGPs stained for EGFP (green), PAX6 (red), and TUJ1 (white), and with DAPI (blue): two RGPs (R-R), one RGP and one postmitotic neuron (R-N), and two postmitotic neurons (N-N). Scale bar, 10 μm. **(H)** Quantification of the percentage of R-R, R-N, and N-N daughter-cell pairs (WT, n = 3 independent replicates; *Nfia;Nfix* cDKO, n = 3 independent replicates). Data are presented as mean ± SEM. *, P < 0.05; **, P < 0.01; n.s., not significant. Statistical analysis was performed using unpaired Student’s t-test.

**Figure S4.**
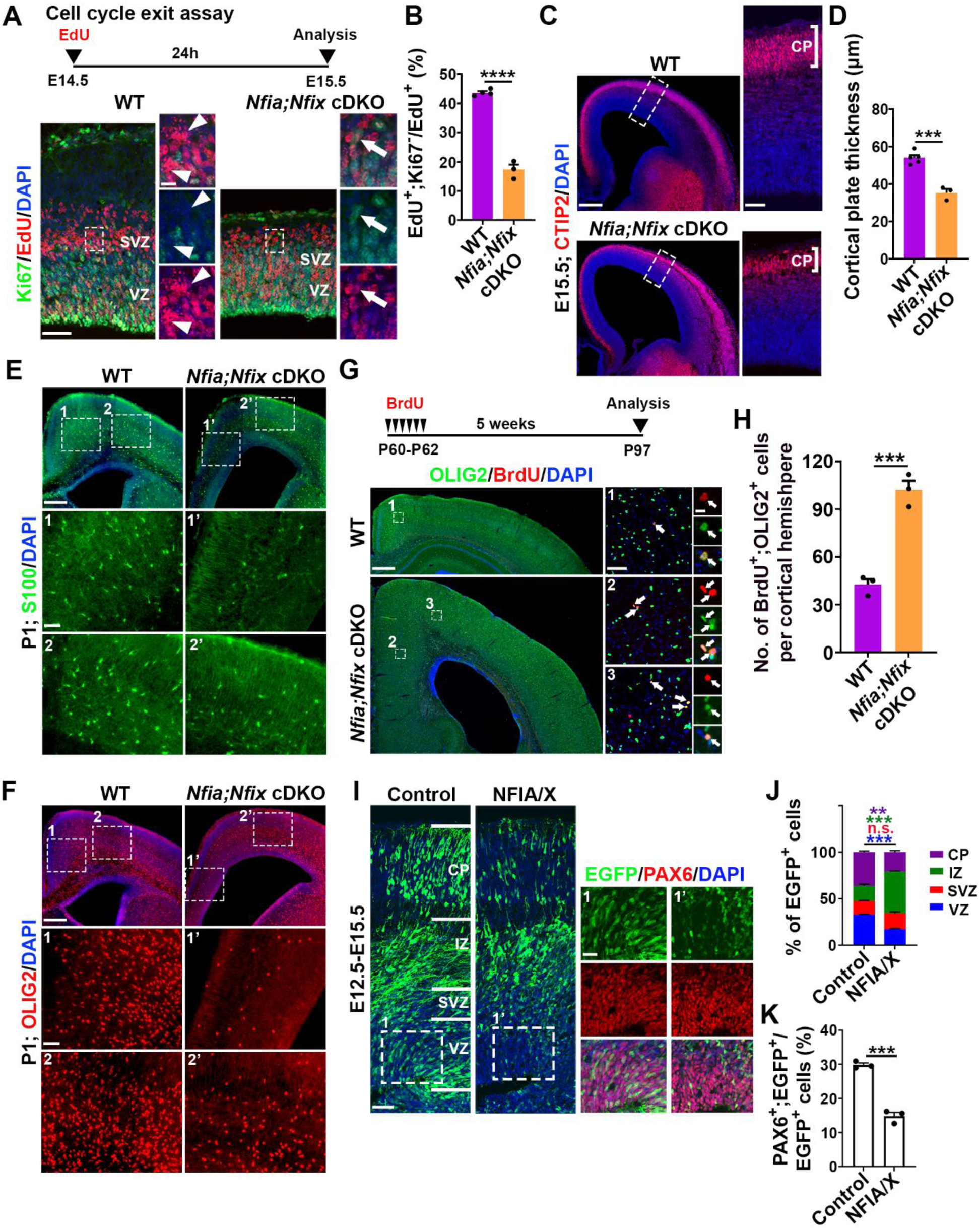
The removal of NFIA and NFIX in RGPs leads to delayed and prolonged neurogenesis and gliogenesis, related to Figure 4 and 5. **(A)** (top) Schematic of cell cycle exit assay. (bottom) Representative images of E15.5 WT and *Nfia;Nfix* cDKO cortices stained for Ki67 (green) and EdU (red), and with DAPI (blue). Arrowheads indicate EdU^+^;Ki67^-^ cells that exited the cell cycle and arrows indicate EdU^+^;Ki67^+^ cells that remain in the cell cycle. Scale bars, 50 μm (left) and 10 μm (right). **(B)** Quantification of the percentage of EdU^+^;Ki67^−^ cells out of the total EdU^+^ cells (WT, n = 4 brains; *Nfia;Nfix* cDKO, n = 3 brains). **(C)** Representative images of E15.5 WT and *Nfia;Nfix* cDKO cortices stained for CTIP2 (red) and with DAPI (blue). High-magnification images of the dorsal cortex are shown to the right. Note the thinner cortical plate (CP) in the *Nfia;Nfix* cDKO brain compared with the WT brain. Scale bars, 250 µm (left) and 50 µm (right). **(D)** Quantification of the thickness of the CP at E15.5 (WT, n = 5 brains; *Nfia;Nfix* cDKO, n = 3 brains). **(E)** Representative images of P1 WT and *Nfia;Nfix* cDKO cortices stained for S100 (green) and with DAPI (blue). High-magnification images (dashed line rectangles) are shown at the bottom. Scale bars, 250 μm (top) and 50 μm (bottom). **(F)** Representative images of P1 WT and *Nfia;Nfix* cDKO cortices stained for OLIG2 (red) and with DAPI (blue). High-magnification images (dashed line rectangles) are shown at the bottom. Scale bars, 250 μm (top) and 50 μm (bottom). **(G)** (top) Schematic of BrdU birthdating assay. (bottom) Representative images of P97 WT and *Nfia;Nfix* cDKO cortices stained for OLIG2 (green) and BrdU (red) administered at P60-P62, and with DAPI (blue). High-magnification images of the medial cortex (dashed line rectangles) are shown to the right. Arrows indicate BrdU^+^;OLIG2^+^ cells. Scale bars, 500 µm (left), 50 µm (middle), and 10 µm (right). **(H)** Quantification of the number of BrdU^+^;OLIG2^+^ cells per cortical hemisphere in WT and *Nfia;Nfix* cDKO cortices (WT, n = 3 brains; *Nfia;Nfix* cDKO, n = 3 brains). **(I)** Representative images of E15.5 cortices electroporated with EGFP control or NFIA and NFIX (NFIA/X) overexpression plasmids together with EGFP (green) at E12.5 stained for PAX6 (red) and with DAPI (blue). High-magnification images of the VZ (dashed line rectangles) are shown to the right. VZ, ventricular zone; SVZ, subventricular zone; IZ, intermediate zone; CP, cortical plate. Scale bars, 50 µm (left) and 25 µm (right). **(J)** Quantification of the percentage of EGFP^+^ cells in different regions of the developing cortex (Control, n = 3 brains; NFIA/X, n = 3 brains). **(K)** Quantification of the percentage of EGFP^+^ cells expressing PAX6 among the total EGFP^+^ cells (Control, n = 3 brains; NFIA/X, n = 3 brains). Data are presented as mean ± SEM. **, P < 0.01; ***, P < 0.001; ****, P < 0.0001; n.s., not significant. Statistical analysis was performed using unpaired Student’s t-test.

**Figure S5.**
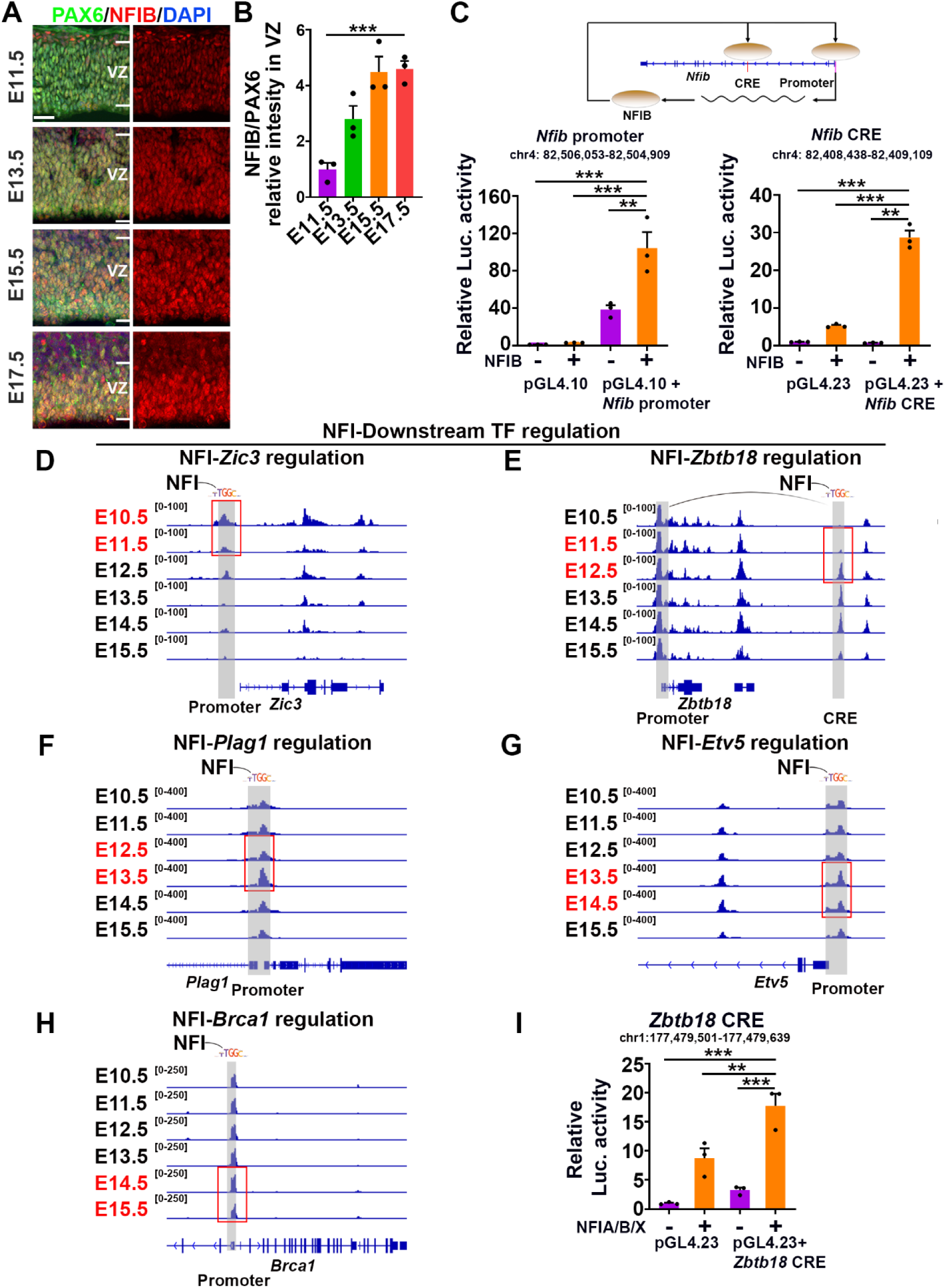
NFIs directly bind to the promoters and/or cis-regulatory elements (CREs) of distinct downstream TFs in RGPs at different developmental stages, related to Figures 6 and 7. **(A)** Representative images of the VZ of the developing mouse neocortices at different embryonic stages (E11.5, E13.5, E15.5, and E17.5) stained for PAX6 (green) and NFIB (red), and with DAP (blue). Note that the expression of NFIB progressively increases in RGPs as development proceeds. White lines indicate the boundaries of VZ. Scale bar, 25 µm. **(B)** Quantification of the relative intensity of NFIB in RGPs compared to that of PAX6. Fold change of expression was calculated by normalizing to the mean level of expression at E11.5 (n = 3 brains for each time point). **(C)** Luciferase reporter assay was performed to measure activity of NFIB-activated transcription from the *Nfib* promoter and CRE. The schematic (top) showed that NFIB protein bound the promoter and CRE of *Nfib* gene and amplified its gene transcription by feed-back positive autoregulation. Co-transfection of the NFIB expression construct and the *Nfib* promoter (left)/*Nfib* CRE (right) resulted in a significantly increased level of relative luciferase activity. n=3 biological replicates per group. Luc, luciferase. **(D)** Representative genome tracks showing accessibility of NFI binding motifs in the promoter (shown in gray box) of E10.5-E11.5 downstream TF *Zic3* at different developmental stages. Note the dynamic accessibility at E10.5-E11.5 transition (highlighted in red rectangle). **(E)** Representative genome tracks showing accessibility of NFI binding motifs in the promoter and CRE (shown in gray boxes) of E11.5-E12.5 downstream TF *Zbtb18* at different developmental stages. Note the dynamic accessibility at E11.5-E12.5 transition (highlighted in red rectangle). **(F)** Representative genome tracks showing accessibility of NFI binding motifs in the promoter (shown in gray box) of E12.5-E13.5 downstream TF *Plag1* at different developmental stages. Note the dynamic accessibility at E12.5-E13.5 transition (highlighted in red rectangle). **(G)** Representative genome tracks showing accessibility of NFI binding motifs in the promoter (shown in gray box) of E13.5-E14.5 downstream TF *Etv5* at different developmental stages. Note the dynamic accessibility at E13.5-E14.5 transition (highlighted in red rectangle). **(H)** Representative genome tracks showing accessibility of NFI binding motifs in the promoter (shown in gray box) of E14.5-E15.5 downstream TF *Brca1* at different developmental stages. Note the dynamic accessibility at E14.5-E15.5 transition (highlighted in red rectangle). **(I)** Luciferase reporter assay was performed to measure activity of NFIA/B/X-regulated transcription from the downstream TF *Zbtb18* cis-regulatory element (CRE). Co-transfection of the NFIA/B/X expression constructs and the *Zbtb18* CRE resulted in a significantly increased level of relative luciferase activity. n=3 biological replicates per group. Luc., luciferase. Data are presented as mean ± SEM. **, P < 0.01; ***, P < 0.001. Statistical analysis was performed using One-way ANOVA.

**Figure S6.**
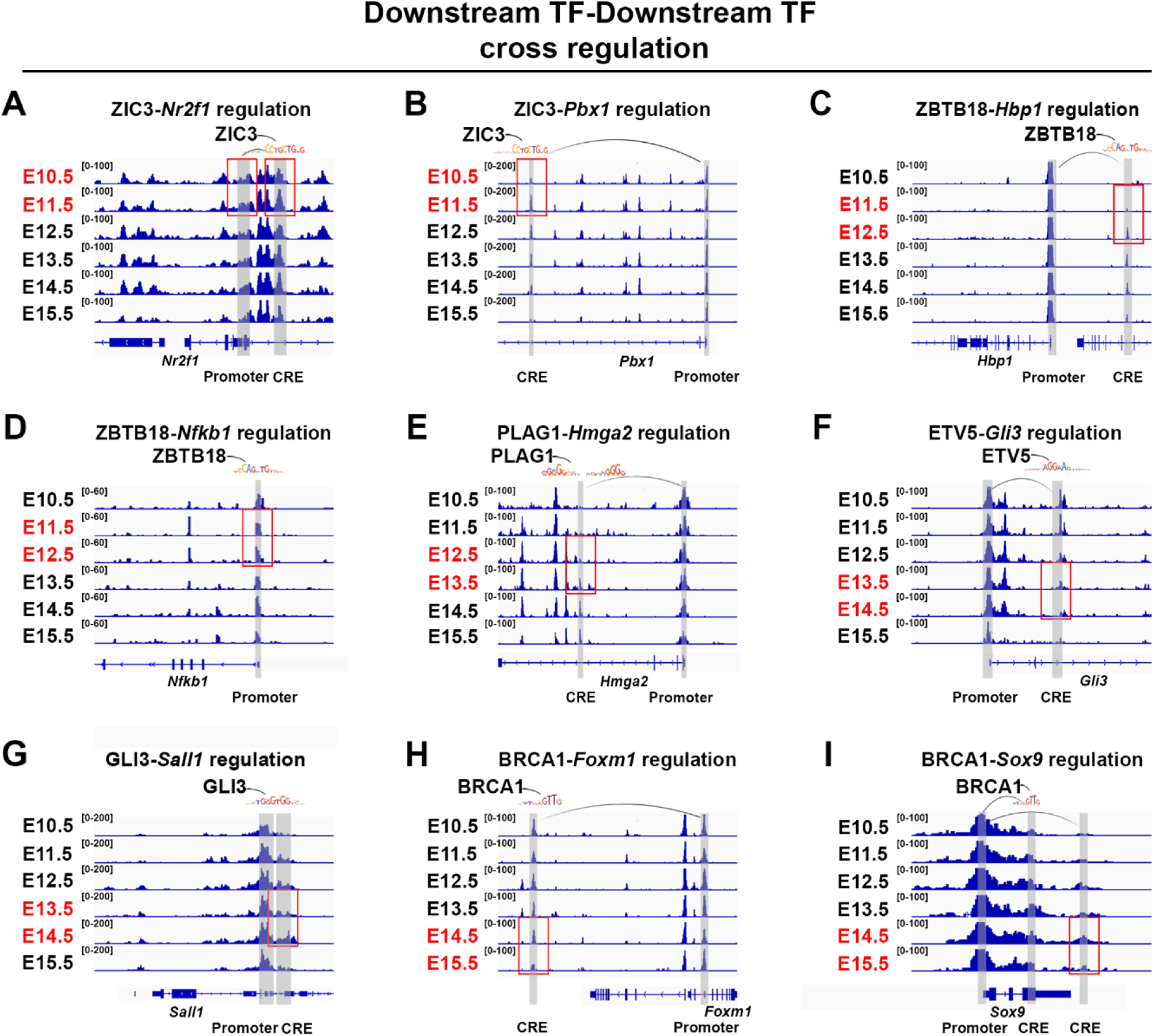
Downstream TF’s cross regulation in RGPs at different developmental stages, related to Figure 7. (**A**) Representative genome tracks showing accessibility of ZIC3 (an E10.5-E11.5 downstream TF) binding motif in the promoter and CRE (shown in gray boxes) of E10.5-E11.5 downstream TF *Nr2f1* at different developmental stages. Note the dynamic accessibility at E10.5-E11.5 transition (highlighted in red rectangles). **(B)** Representative genome tracks showing accessibility of ZIC3 (an E10.5-E11.5 downstream TF) binding motif in the promoter and CRE (shown in gray boxes) of E10.5-E11.5 downstream TF *Pbx1* at different developmental stages. Note the dynamic accessibility at E10.5-E11.5 transition (highlighted in red rectangle). **(C)** Representative genome tracks showing accessibility of ZBTB18 (an E11.5-E12.5 downstream TF) binding motif in the promoter and CRE (shown in gray boxes) of E11.5-E12.5 downstream TF *Hbp1* at different developmental stages. Note the dynamic accessibility at E11.5-E12.5 transition (highlighted in red rectangle). **(D)** Representative genome tracks showing accessibility of ZBTB18 (an E11.5-E12.5 downstream TF) binding motif in the promoter (shown in gray box) of E11.5-E12.5 downstream TF *Nfkb1* at different developmental stages. Note the dynamic accessibility at E11.5-E12.5 transition (highlighted in red rectangle). **(E)** Representative genome tracks showing accessibility of PLAG1 (an E12.5-E13.5 downstream TF) binding motif in the promoter and CRE (shown in gray boxes) of E12.5-E13.5 downstream TF *Hmga2* at different developmental stages. Note the dynamic accessibility at E12.5-E13.5 transition (highlighted in red rectangle). **(F)** Representative genome tracks showing accessibility of ETV5 (an E13.5-E14.5 downstream TF) binding motif in the promoter and CRE (shown in gray boxes) of E13.5-E14.5 downstream TF *Gli3* at different developmental stages. Note the dynamic accessibility at E13.5-E14.5 transition (highlighted in red rectangle). **(G)** Representative genome tracks showing accessibility of GLI3 (an E13.5-E14.5 downstream TF) binding motif in the promoter and CRE (shown in gray boxes) of E13.5-E14.5 downstream TF *Sall1* at different developmental stages. Note the dynamic accessibility at E13.5-E14.5 transition (highlighted in red rectangle). **(H)** Representative genome tracks showing accessibility of BRCA1 (an E14.5-E15.5 downstream TF) binding motif in the promoter and CRE (shown in gray boxes) of E14.5-E15.5 downstream TF *Foxm1* at different developmental stages. Note the dynamic accessibility at E14.5-E15.5 transition (highlighted in red rectangle). **(I)** Representative genome tracks showing accessibility of BRCA1 (an E14.5-E15.5 downstream TF) binding motif in the promoter and CREs (shown in gray boxes) of E14.5-E15.5 downstream TF *Sox9* at different developmental stages. Note the dynamic accessibility at E14.5-E15.5 transition (highlighted in red rectangle).

**Figure S7.**
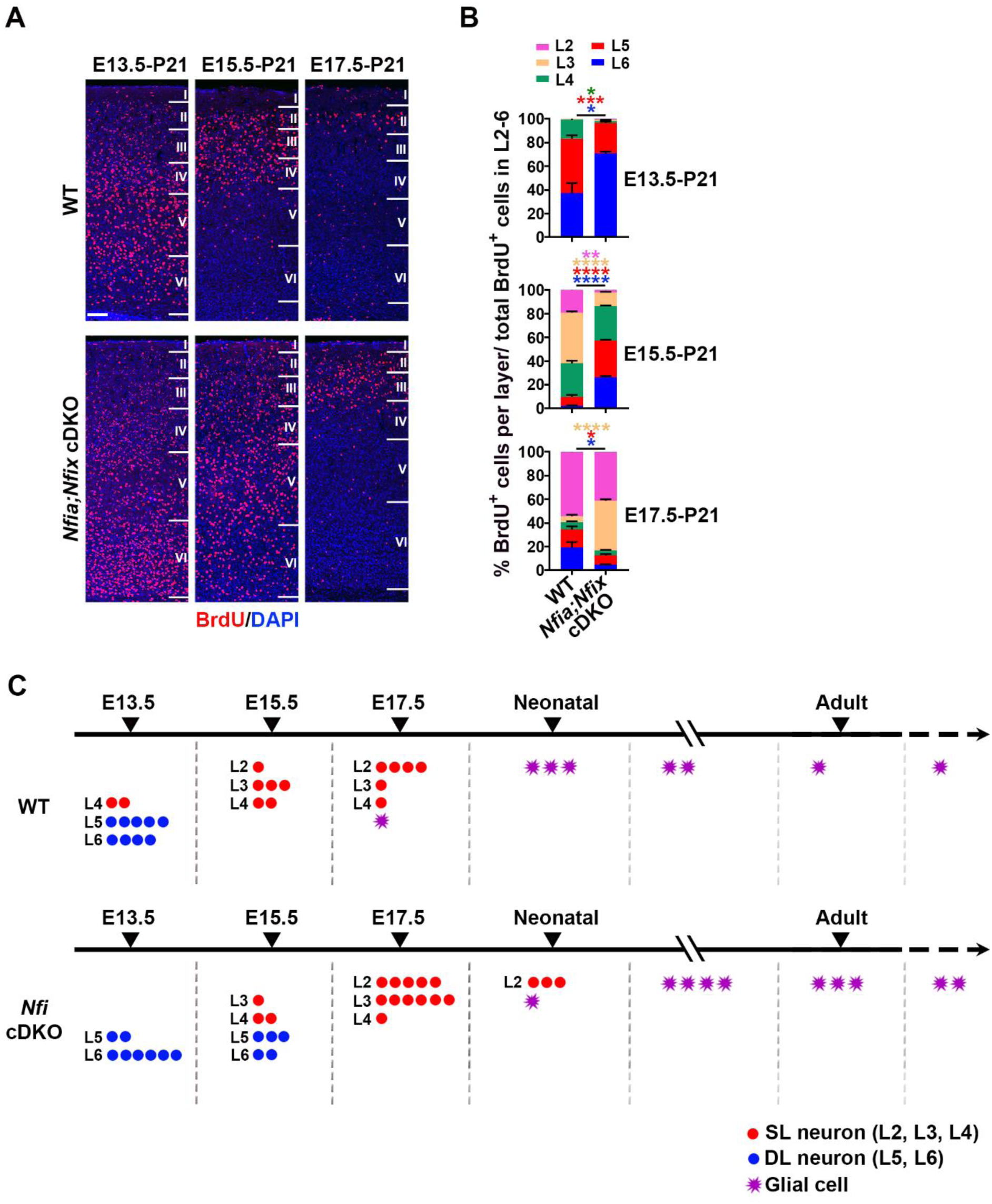
The removal of NFIA and NFIX in RGPs shifts neuronal progeny output toward early developmental stages, related to Figure 8. **(A)** Representative images of P21 WT and *Nfia;Nfix* cDKO cortices stained for BrdU (red) injected at E13.5 (left), E15.5 (middle), and E17.5 (right), and with DAPI (blue). Scale bar: 100 µm. **(B)** Quantification of the percentage of BrdU^+^ cells in different layers of the cortex (E13.5: WT, n = 4 brains; *Nfia;Nfix* cDKO, n = 3 brains; E15.5: WT, n = 3 brains; *Nfia;Nfix* cDKO, n = 4 brains; E17.5: WT, n = 4 brains; *Nfia;Nfix* cDKO, n = 4 brains). **(C)** Schematic showing loss of NFIA and NFIX protracts RGP lineage progression in the cortex. DL, deep layer; SL, superficial layer. Data are presented as mean ± SEM. *, P<0.05; **, P < 0.01; ***, P < 0.001; ****, P < 0.0001. Statistical analysis was performed using unpaired Student’s t-test.

**Figure S8.**
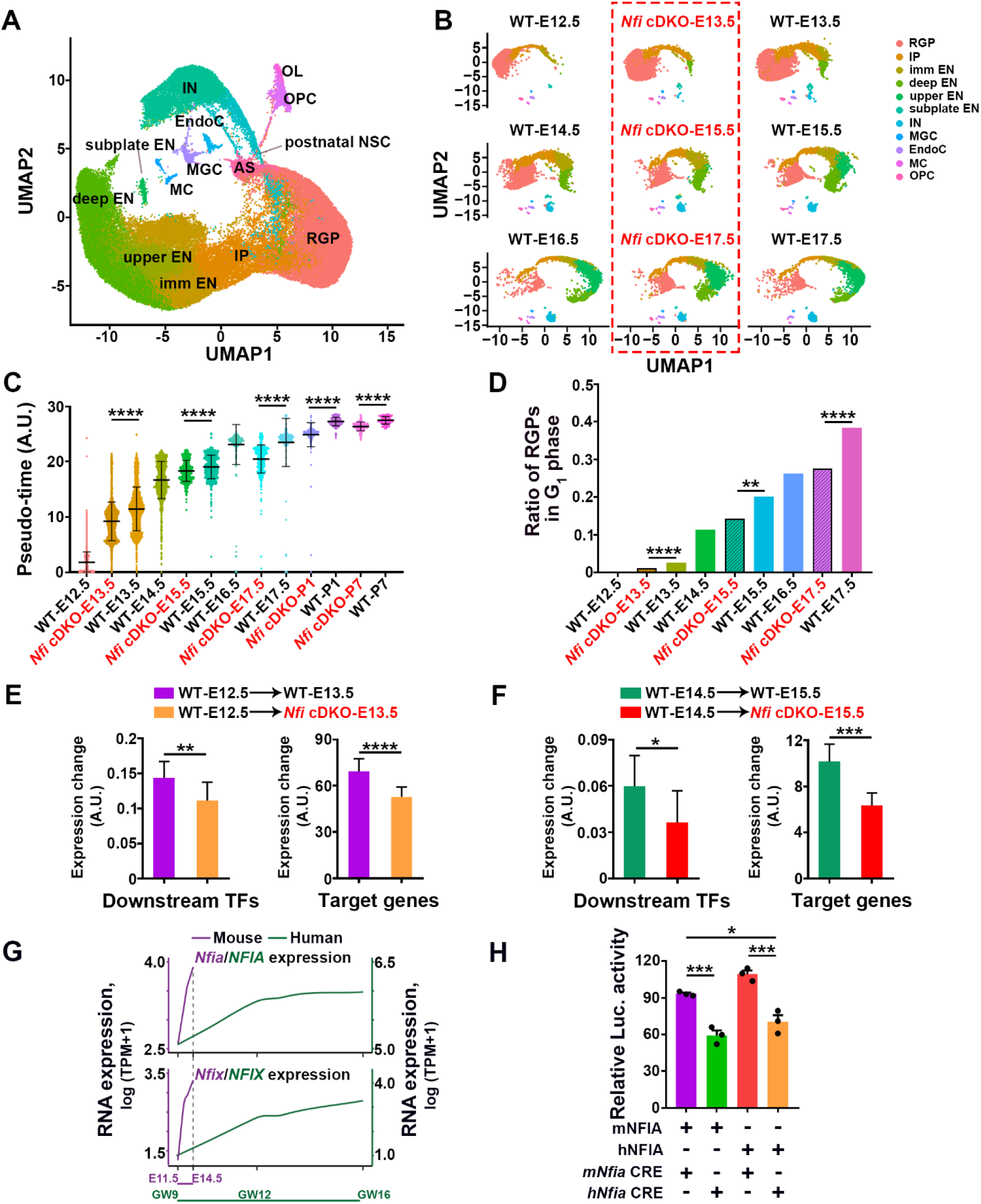
The removal of NFIA and NFIX in RGPs leads to younger temporal identity, related to Figure 8. (A) UMAP-dimension reduction of the expression profiles of individual cells obtained in WT and *Nfia;Nfix* cDKO cortices at E12.5-P7. RGP, radial glial progenitor; IP, intermediate progenitor; imm EN, immature excitatory neuron; deep EN, deep layer excitatory neuron; upper EN, upper layer excitatory neuron; subplate EN, subplate excitatory neuron; IN, interneuron; OPC, oligodendrocyte precursor cell; EndoC, endothelial cell; MGC, microglial cell; MC, meningeal cell; postnatal NSC, postnatal neural stem cell; AS, astrocyte. (B) UMAP-dimension reduction of the expression profiles of individual cells obtained in WT and *Nfia;Nfix* cDKO (*Nfi* cDKO) cortices at different developmental stages (E12.5-E17.5). RGP, radial glial progenitor; IP, intermediate progenitor; imm EN, immature excitatory neuron; deep EN, deep layer excitatory neuron; upper EN, upper layer excitatory neuron; subplate EN; subplate excitatory neuron; IN, interneuron; MGC, microglial cell; EndoC, endothelial cell; MC, meningeal cell; OPC, oligodendrocyte precursor cell. (C) Developmental progression score analysis of the temporal identity of WT and *Nfia;Nfix* cDKO RGPs. (D) Quantification of the ratio of RGPs in G_1_ phase of WT and *Nfia;Nfix* cDKO (*Nfi* cDKO) cortices across different embryonic stages. (E) Quantification showing a reduction in the magnitude of expression changes of downstream TFs and their target genes in *Nfia;Nfix* cDKO RGPs compared with WT control RGPs during E12.5-E13.5 transition. (F) Quantification showing a reduction in the magnitude of expression changes of downstream TFs and their target genes in *Nfia;Nfix* cDKO RGPs compared with WT control RGPs during E14.5-E15.5 transition. (G) Progressive dynamics of *Nfia*/*NFIA* (top) and *Nfix*/*NFIX* (bottom) expression in RGPs along aligned cortical development based on mouse (purple) or human (green) single-cell RNA-seq data (GSE104276) (*99*). Embryonic day (E); Gestational week (GW). (H) Luciferase assay using the *Nfia* (mouse) or *NFIA* (human) CRE-Luc reporter co-transfected with the indicated amounts of the same or different species-mouse or human NFIA expression plasmids (pCIG-NFIA) or control plasmids (pCAG-IRES-EGFP). Fold change of relative Luc activity was calculated by normalizing to the mean Luc activity level of empty Luc reporter co-transfected with pCIG plasmid control group. (n = 3 biological replicates for each group). Luc, luciferase. Data are presented as mean ± SD (**C)** and mean ± SEM (**D-F and H)**. *, P < 0.05; **, P < 0.01; ***, P < 0.001; ****, P < 0.0001. Statistical analysis was performed using unpaired Student’s t-test **(C)**, Chi-square test **(D)**, and pairwise Wilcoxon rank sum test **(E and F)**, and One-way ANOVA **(H)**.

